# Perturbation of ACE2 structural ensembles by SARS-CoV-2 spike protein binding

**DOI:** 10.1101/2021.03.02.433608

**Authors:** Arzu Uyar, Alex Dickson

## Abstract

The human ACE2 enzyme serves as a critical first recognition point of coronaviruses, including SARS-CoV-2. In particular, the extracellular domain of ACE2 interacts directly with the S1 tailspike protein of the SARS-CoV-2 virion through a broad protein-protein interface. Although this interaction has been characterized by X-ray crystallography and Cryo-EM, these structures do not reveal significant differences in ACE2 structure upon S1 protein binding. In this work, using several all-atom molecular dynamics simulations, we show persistent differences in ACE2 structure upon binding. These differences are determined with the Linear Discriminant Analysis (LDA) machine learning method and validated using independent training and testing datasets, including long trajectories generated by D. E. Shaw Research on the Anton 2 supercomputer. In addition, long trajectories for 78 potent ACE2-binding compounds, also generated by D. E. Shaw Research, were projected onto the LDA classification vector in order to determine whether the ligand-bound ACE2 structures were compatible with S1 protein binding. This allows us to predict which compounds are “apo-like” vs “complex-like”, as well as to pinpoint long-range ligand-induced allosteric changes of ACE2 structure.

## Introduction

Human angiotensin-converting enzyme 2 (ACE2) is a metallo-carboxypeptidase that cleaves isoforms I and II of the angiotensin peptide. ACE2 is a key regulator of blood volume and is expressed in tissues throughout the cardiovascular system, in arterial smooth muscle cells, and endothelial cells in small and large arteries. Together with the proto-oncogene MAS, ACE2 also functions to reduce acute injury and inhibit fibrogenesis in the lungs.^1^ Structurally, ACE2 is a zinc metallopeptidase, 805 residues in length. It has three domains: extracellular (residues 18-740), transmembrane (residues 741-761) and cytoplasmic (residues 762-805) (Fig. 1A). Notably, ACE2 acts as the host receptor for viral entry of coronaviruses, including severe acute respiratory syndrome coronavirus 2 (SARS-CoV-2),^2,3^ which is responsible for the coronavirus disease (COVID-19) pandemic. Two other coronaviruses, namely SARS-CoV^4^ and HCoV-NL63,^5^ also use ACE2 as a primary means of viral entry. All three of these coronaviruses use the S1 subunit of their extra-cellular spike glycoproteins to bind to ACE2 as a host receptor. Therefore, a better understanding of how to prevent ACE2-S1 interactions could be a key component in drug development efforts to combat coronavirus infection.

**Figure 1:**
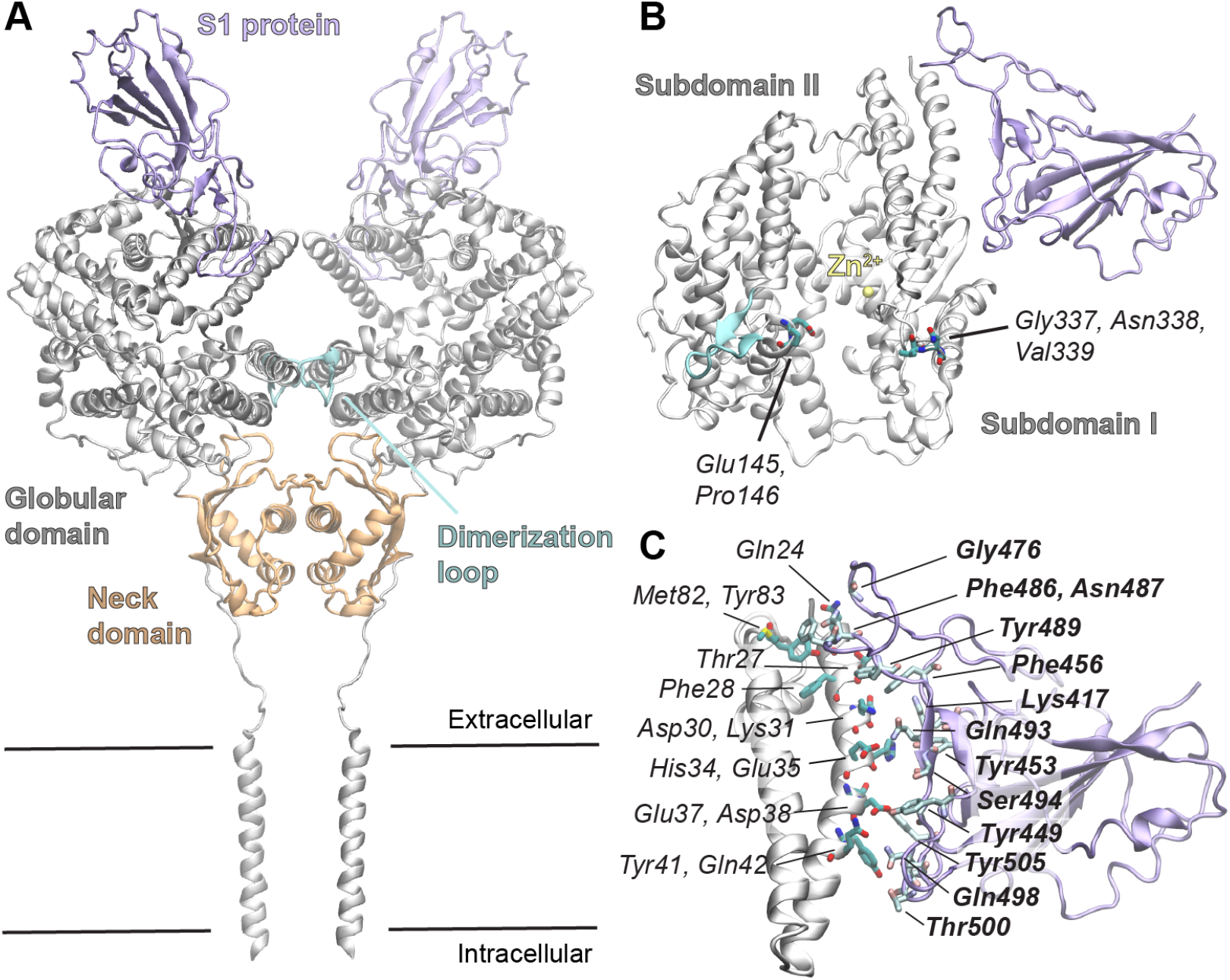
Overview of ACE2 structure. (A) A view of the ACE2 dimer structure interacting with the viral S1 protein (PDB: 6M17^6^). (B) The extracellular peptidase domain of ACE2 and the S1 protein. Key residues discussed in the text are labeled and shown in licorice representation. The catalytic Zn^2^+ ion is shown in yellow. (C) A view of the molecular interactions between helices 1 and 2 of the peptidase domain with the S1 protein. Specific interactions were determined using the PDBSum database^7,8^ and interacting residues are labeled and shown in licorice representation. Residues from the S1 protein are shown in bold.

The extracellular part of ACE2 is composed of a small neck domain and a large clam-shell-shaped globular domain (also referred to as the “peptidase domain”) that is divided by a deep channel, approximately 60 Å in length, into subdomains I and II (Fig. 1B). Subdomain I involves the active site of the enzyme with a bound zinc ion as a cofactor (yellow sphere in Fig. 1B). The SARS-CoV-2 S1 protein attaches to the N-terminal helix (H1) in subdomain I. Another functionally important loop region (residues 131 to 142) is located on subdomain II, which is involved in ACE2 dimerization with the neck domain (residues 616 to 726).^6^ ACE2 has seven N-linked glycosylation sites although none of these play a major role in S1 protein binding. However, there is recent evidence that some glycosylated residues can form direct hydrogen bonding interactions with S1^9–11^ and in particular that Asn90 plays a stabilizing role in ACE2-S1 interactions,^12^ although this evidence is mixed.^13^

The details of ACE2 interaction with the S1 proteins of SARS-CoV-2 and of other ACE2-hosted coronaviruses are of great interest for the discovery of receptor-virus interaction inhibitors. Several MD simulation and modeling studies recently showed that SARS-CoV-2 binds to ACE2 stronger than other coronaviruses and identified critical interacting residues^14–20^ and suggested key mutations in the conserved residues on S1 spike protein that would lower binding affinity.^21^ Hadi-Alijanvand and Rouhani^22^ determined mutations on ACE2 that affect affinity between ACE2 and S1 protein using several fast computational predictors, such as PISA^23^ and FoldX.^24^ They proposed mutations that enhances affinity and found one mutation (V485L) that causes a lower affinity than the wild-type ACE2. They also proposed closed conformations of ACE2 have higher affinity to S1 than that of open state conformations.

The extracellular domain of ACE2 is homologous to other metallo-peptidases such as ACE,^25^ carboxypeptidase Pfu^26^ and neurolysin,^27^ which all share the characteristic “clam-shell” structure. Neurolysin dynamics were studied in detail in our previous work^28^ using MD simulations and the Linear Discriminant Analysis (LDA) machine learning method. Our findings compared contact networks of the apo neurolysin and neurolysin bound to an allosteric inhibitor and pinpointed several differences that were far from inhibitor binding site, which suggests long-range allosteric behavior in metallo-peptidases.

Motivated by the devastating impact of the COVID-19 epidemic, the dynamics of ACE2 homodimers has been recently investigated in a number of studies using molecular dynamics simulation. Mohammad et al.^29^ simulated wild-type homodimer ACE2 and its N720D mutant form to compare the affinity of ACE2 to type-II transmembrane serine protease (TMPRSS2) that cleaves ACE2 on its N-terminus, which is required for the interaction of ACE2 with the S protein. Their binding affinity predictions and MD simulations showed that the mutant form has higher affinity, therefore they propose that N720D mutant ACE2 might be more susceptible to S1 protein binding. Another study by Barros et al.^11^ takes into account the membrane in their 1.0 *µ*s simulation for the apo and complex ACE2 with the spike protein. They compared the head tilt angles of apo and complex ACE2 from the membrane and showed that, due to ACE2 flexibility, more than one ACE2 protein can be accommodated by a single spike protein in the up conformation. Earlier in 2020, a series of long MD simulations for the apo and S1-bound ACE2 were performed on the Anton 2 supercomputer^30^ and made available to download.^31^ They also examined 5152 molecules in an FDA-investigational drug library^32^ using MD simulations and docking, and selected 78 out of these molecules that remained bound to ACE2. These candidate compounds are located at three distinct regions on ACE2 that are involved in either interaction with spike^6,9^ or ACE2 homo-dimerization.^6^

Here, we analyze recently obtained crystal structures of both the apo and SARS-CoV-2 S1 protein-bound structures of the ACE2 peptidase domain, to identify differences between these two forms, searching for the hallmarks of S1 protein binding in the ACE2 structure. Differences in these structural ensembles has the potential to inform the design of allosteric inhibitors of ACE2-S1 interactions that slow or prevent viral entry. We then perform several independent all-atom MD simulations using CHARMM^33–35^ for the apo and spike S1 protein bound forms of the ACE2 peptidase domain, and compare differences observed in these simulations with those in the ACE2 crystal structures. We further apply LDA to automatically detect structural differences between apo and complex ACE2 and generate several LDA classification vectors that can quantify ACE2 conformations on a spectrum of “apo-like” versus “complex-like”. These vectors are then applied to a massive simulation dataset of 78 ACE2-binding ligands recently made available be D.E. Shaw Research, allowing us to classify each of these compounds as “apo-generating” or “complex-generating” based on their structural impact on the ACE2 peptidase domain. We hope these findings will illuminate critical sites for allosteric modulation of ACE2-S1 interactions, as well as provide a general study of the relationship between ligand binding and conformational change in metallo-peptidases.

## Results

### Comparison of apo and S1-bound ACE2 crystal structures

As a starting point we first examine differences that can be gleaned from the apo and S1-bound ACE2 crystal structures alone. This reveals little information as the structures show only slight differences: alignment of the apo (PDB:1R42^36^) and the S1-bound, referred to hereon as “complex”, (PDB:6M17^6^) ACE2 crystal structures results in a C*α* RMSD of 0.78 Å. To determine the regions of ACE2 that differ most significantly between the structures, we calculate the C*α* RMSD for each residue. The highest differences (above a cutoff of 0.20 Å) are found in the N-terminus (Ile21), the central loop (residues 331-347) in subdomain I, and Leu424, Pro426, Asp427, and Phe428 on the bottom loop of the subdomain I. Residue-level RMSD was also calculated between the structures after thermal equilibration and very similar results were obtained.

The S1 protein binding interface involves the N-terminus helices (H1 and H2: residues 21-100) of ACE2. Although this region is directly perturbed by the S1 protein, we do not observe significant differences (aside from Ile21) in the protein backbone. Using all-atom RMSD we identify differences in side chain conformation for residues Ile21, Glu23, Lys31, Met82 and Gln89 that exceed a cutoff of 0.45 Å. Residues in this region that are interacting with S1 are shown in Fig. 1C, including Lys31 and Met82. RMSD profiles between the apo and the complex ACE2 structures can be seen in Figure S1.

We go further to compare the backbone dihedral angles (*ϕ, Ψ* and *ω*) between the two crystal structures (Fig. 2, left). The shaded red region identifies residues with significantly different values (> *π/*2) of one or more of these dihedral angles, which are summarized in Table 1. *ϕ* angle comparison between the apo and the complex crystal structures revealed six residues as outliers and Asp213 is the most different one (top outlier), which is on the loop (residues 205-219) at the backside of ACE2. From *Ψ* angle comparison between ACE2 crystal structures, seven residues are revealed as outliers and Gln325 near S1-binding site is the top outlier. The most striking difference was observed from the Glu145/Pro146 *ω* angle, which resulted from the different isomerization states of the proline residue.

**Table 1:** Key residues (outliers) determined from dihedral angle analysis between the apo and the complex structures of ACE2. Asterices indicate outlier residues with the largest differences. Molecular dynamics outliers (MD) are determined using both the most probable angle and the “Earth mover’s distance” (EMD) between the probability distributions.

|  | Crystal structure outliers | MD outliers (most probable) | MD outliers (EMD) |
| --- | --- | --- | --- |
| $\phi$ | Ser106, Asp213*, Gln325, Gly326, Gly337, Val339, His345 | Gly211*, Gly326, Gly337, Val339, Asn394 | Glu145, Gly147, Gly326*, Val339, Asn394 |
| $\psi$ | Ile21, Ser105, Glu145, Thr324, Gln325*, Pro336, Gly337, Val339 | Glu145, Gly211, Ile291*, Gln325, Pro336, Gly337, Val339 | Ile21, Glu145*, Ile291, Gln325, Pro336, Gly337, Val339 |
| $\omega$ | Glu145 | Glu145 | Glu145 |

**Figure 2:**
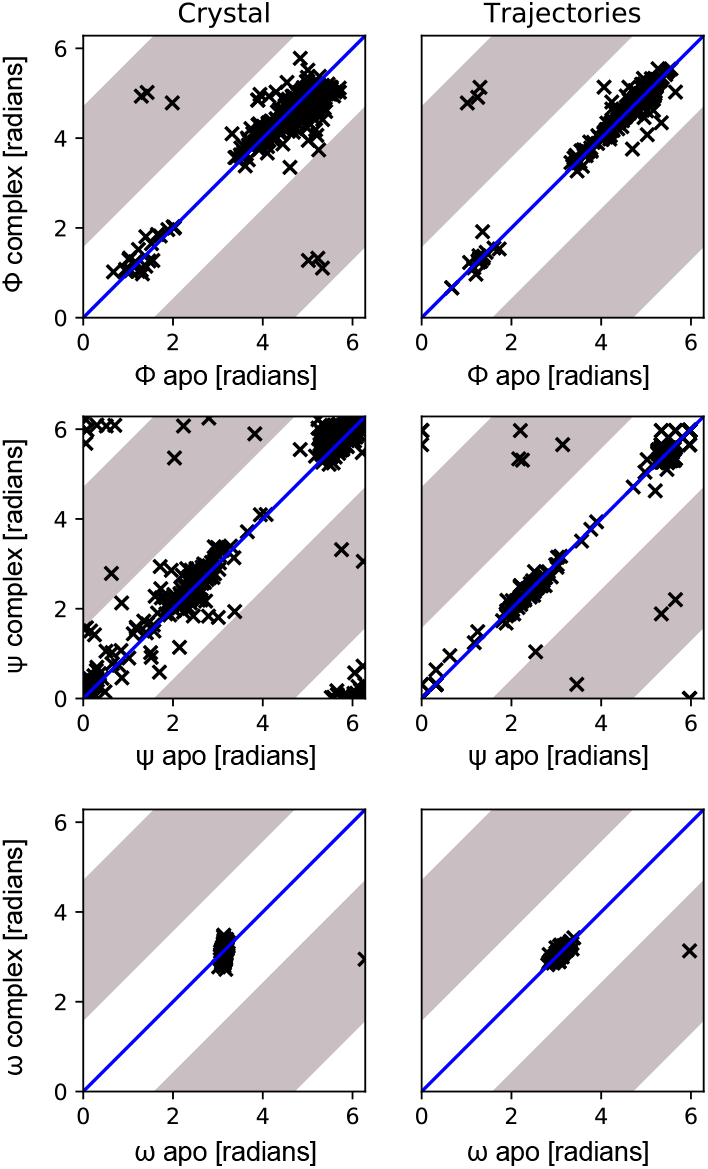
Backbone dihedral angle comparison between apo and complex forms of ACE2. Scatter plots compare the dihedral angles of ACE2 alone (apo) vs ACE2 bound to the S1 protein (complex). Differences obtained between the crystal structures are shown on the left and differences in MD trajectories are shown on the right. Areas with significant differences (greater than ±*π/*2) are shown in grey. Differences are examined separately for dihedral angles *ϕ* (top), *Ψ* (middle) and *ω* (bottom).

Overall, these two methods of structural comparison had little overlap and it is not clear whether these differences between the crystal structures will persist between the apo and complex forms of ACE2 in a biologically relevant environment. We next conducted a series of all-atom molecular dynamics simulations of both the apo and S1-bound systems in order to identify significant structural differences in ACE2 that occur upon S1-protein binding.

### MD simulations reveal differences in critical dihedral angles in ACE2

We simulated a set of 15 trajectories for each of the apo-ACE2 and ACE2-S1 systems, each one 100 ns in length. Throughout the simulations for the complex, the ACE2-S1 interface remained intact. We repeat the *ϕ, Ψ* and *ω* analysis, this time by computing probability distributions for each angle. Fig. 2 (right side) compares the most probable angles of these distributions between the apo and S1-bound forms of ACE2. Outliers are again identified using differences > *π/*2 and summarized in Table 1. The MD outliers showed significant overlap with those computed directly from the crystal structures, although some unique information was obtained. For *ϕ* three of six of the crystal structure differences persisted during the MD trajectories. For *Ψ* four of seven of the crystal structure differences persisted, although in addition the MD trajectories identified three new residues (Gly211, Ile291, Val339) where differences were not found in the crystal structures. The comparison of *ω* angles again showed that Glu145/Pro146 is the only clear significant difference. As expected we did not observe any proline isomerization changes in either the apo or complex trajectories.

Unfortunately, when angle probability distributions have multiple peaks, the above measurement can be sensitive to small changes that affect which angle is the global maximum. To address this we also determine outlier residues using the “Earth-mover’s distance” between the probability distributions of each backbone angle, which essentially measures how much total probability needs to be moved in order to transform one probability distribution into the other. This was achieved using the pyemd package,^37,38^ while taking the periodic nature of the angles into account. Again we observe moderate overlap with the crystal structure outliers, although there were changes in the angles that were considered the most significantly different.

In summary, some of the differences in crystal structure backbone angles between the apo and the complex crystal structures are also observed in probability distributions obtained from the simulations. Differences in the central loop of subdomain I (Gly337, Asn338, Val339) show up persistently in the backbone analysis as well as the residue-level RMSD analysis. Most clearly, the *ω* angle analysis clearly points to a single residue that is observed both in the crystal structures and simulations. While this type of structural analysis can be useful to point us to specific residues and regions of the protein, it does not provide a way to synthesize this information. For instance, if we were presented with another ACE2 structure and asked to predict whether it is compatible with the apo or S1-bound form, we could look individually at central loop conformations, or specific backbone angles, but there is no clear way to combine this information into a single number that can be used for scoring. Next we employ the LDA method to do exactly that.

### LDA automatically detects ACE2 structural differences

We applied LDA to the apo and the complex trajectories (15 runs for each) to identify structural differences between the two forms of ACE2. LDA yields a classification vector – analogous to collective variables in principal component analysis (PCA) – that results in the largest differences between the classes, while maintaining small differences within the classes.^39,40^ As here there are only two classes during training (“apo” and “complex”), the analysis yields only one LDA vector. Projection of a structure onto this vector yields a single scalar, which in this case assesses whether the structure is “apo-like” or “complex-like”. We split the apo and the complex data into three subsets of five runs each in order to evaluate both training and testing accuracy as summarized in Table 2. To rigorously test the generalizability of the LDA vectors, we train on one third of the data and test on the remaining two thirds. Accuracy values for each state (“apo” or “complex”) are calculated as the fraction of samples that are successfully predicted in that state, using the LDA projection value and a cutoff distinguishing the two states.

**Table 2:** Training-testing accuracy table for LDA method. “DR” denotes that the DESRES dataset was included in the training.

| Predictor | $n_{\text{atoms}}$ | $n_{\text{train}}$ | training accuracy | | testing accuracy | |
| --- | --- | --- | --- | --- | --- | --- |
|  |  |  | apo | complex | apo | complex |
| CA-592 | 592 | 5 | $1.000 \pm 0.000$ | $1.000 \pm 0.000$ | $0.990 \pm 0.008$ | $0.992 \pm 0.007$ |
| CA-592 | 592 | 15 | 1.000 | 1.000 | - | - |
| CA-590 | 590 | 5 | $1.000 \pm 0.000$ | $1.000 \pm 0.000$ | $0.966 \pm 0.024$ | $0.966 \pm 0.022$ |
| CA-590 | 590 | 15 | 1.000 | 1.000 | - | - |
| CA-592-DR | 592 | 5 | $1.000 \pm 0.000$ | $1.000 \pm 0.000$ | $0.9813 \pm 0.014$ | $0.976 \pm 0.024$ |
| CA-592-DR | 592 | 15 | 1.000 | 1.000 | - | - |
| CA-INT | 80 | 5 | $0.88 \pm 0.10$ | $0.93 \pm 0.02$ | $0.83 \pm 0.09$ | $0.74 \pm 0.20$ |
| CA-INT | 80 | 15 | 0.85 | 0.89 | - | - |
| CA/SC-INT | 158 | 5 | $0.986 \pm 0.013$ | $0.993 \pm 0.004$ | $0.908 \pm 0.048$ | $0.84 \pm 0.14$ |
| CA/SC-INT | 158 | 15 | 0.968 | 0.982 | - | - |

We first build a predictor using all of the ACE2 C*α* atoms (residues 21-612), denoted as “CA-592”, as we use 592 C*α* atoms in the feature set. Training using each of the three subsets yields a good separation between the apo and the complex data sets and a prediction accuracy value of 1.00 in the training set. The testing accuracy values of this training are only slightly lower compared to the training accuracy values (0.95-1.00). As expected, training using all runs (1-to-15) again gives a prediction accuracy of 1.00, although no data remains for testing. Probability distributions of LDA projections resulting from different feature sets are provided in Figure S2. Predictors with perfect accuracy yield probability distributions that are completely separated. As these distributions start to overlap, particularly with testing data, the accuracy falls below 1.00. Displacements in the LDA mode of CA-592 performed using all runs are shown using a vector representation in Fig. 3 where key residues are denoted. Atomic intensity values for this LDA mode are calculated using the components of the LDA vector (see Methods) and shown in Figure S3. Here, CA-592 automatically predicted that the most significant difference between the apo and complex ACE2 can be explained with two residues: Glu145 and Pro146, which is consistent with the dihedral angle analysis above.

**Figure 3:**
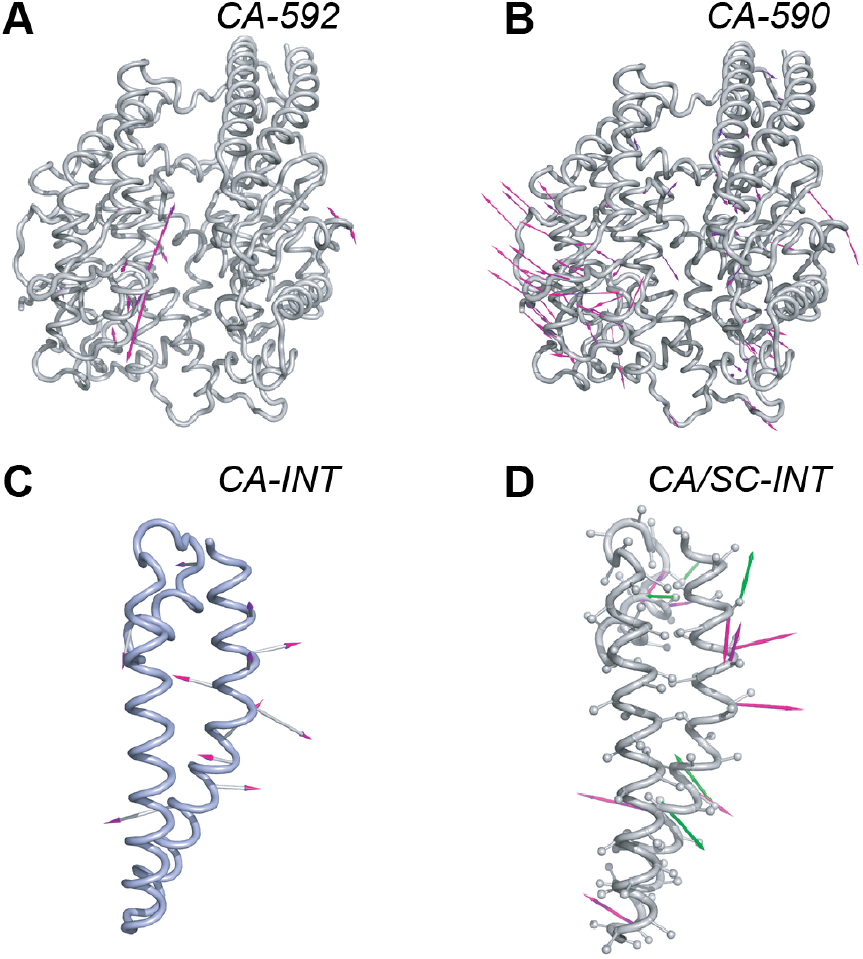
Mode vectors from linear discriminant analysis. Each panel shows LDA vectors visualized as arrows on the ACE2 structure. Figure labels refer to specific LDA vectors obtained with different molecular features, as described in the text. For CA-592, CA-590 and CA-INT, the pink arrows show the contribution of each C*α* atom to the LDA vector. For CA/SC-INT, the pink arrows show the C*α* contributions and the green arrows show contributions from the sidechains. The bottom two structures show only the interface region of ACE2 composed of helices 1 and 2.

As mentioned earlier, D.E. Shaw Research (DESRES) performed 10-microsecond long MD simulations for the apo and complex ACE2^31^ on the Anton 2 supercomputer.^30^ We added these trajectories to our set of 15 apo and complex trajectories and applied LDA to the combined set; this LDA vector is denoted as CA-592-DR. The training and testing accuracy values on our trajectories are slightly lower than that of CA-592, indicating that the conformational differences between the two sets of trajectories impaired the generalizability among the trajectories generated here. However, adding DESRES and our data together yielded a similar LDA mode as that of CA-592 (Figure S3 (B) and (D)). Again, Glu145 and Pro146 appeared as by far the largest components of the CA-592-DR mode. It is worth noting that the DESRES trajectories began in the same isomerization states for Pro146 as those used in our work. Comparison of atomic intensities for CA-592-DR and CA-592 is provided in Figure S3 (C).

The dominance of Glu145 and Pro146 in the LDA modes of CA-592 and CA-592-DR is clear and consistent, but unfortunately the projections will be dominated by the positions of these two residues and other residues that contribute to apo/complex conformational differences will be obscured. We thus removed these two residues from the feature vectors and repeated the LDA for our 15 apo and complex trajectories; this predictor is denoted as CA-590. The training and testing accuracy values for each subset and the whole data set in CA-590 are similar to that of CA-592 (Table 2). In this case, the LDA mode of CA-590 has a more collective behavior that involves the displacements of the bottom half of the subdomain I and II and the bottom part of the N-terminus helices 1 and 2. Some of the significant key residues are on: the dimerization loop (Pro138 and Gln139); on the loop that connects the beta sheets in subdomain I (Pro336, Gln340, and Lys341); and on the loop that is near bottom arms of the S1-protein (Gly326) as highlighted in Fig. 3B. Atomic intensity values for CA-590 and comparison with that of CA-592 are also provided in Figure S3 (E) and (F), respectively. Compared to all the differences obtained from the backbone angle and residue-level RMSD analysis, the LDA analysis revealed 9 new residues.

In addition, we introduce another predictor that focuses on the ACE2-S1 binding interface by only taking into account residues in the first two helices of the N-terminus (residues 21-100) in the LDA calculations. For the binding interface, we performed LDA in two different ways: i) using only CA atoms (named “CA-INT”) and ii) using CA atoms and the centers of geometry of the sidechain heavy atoms (named “CA/SC-INT”, where SC represents “sidechain”). Both training and testing accuracy values for CA-INT are lower than that of CA-590 and CA-592 (Table 2). Fig. 3C shows displacement vectors for CA-INT where 11 key residues are labeled. Seven of them are S1-interacting residues (data taken from PDBsum^7,8^) and are underlined in the figure. On the other hand, CA/SC-INT revealed 10 key residues where three of these are not found by CA-INT (Lys26, Asn53 and Val93) as shown in Fig. 3D. Thr27 and Tyr41 are the only key residues where both CA and SC atoms are revealed as significant and they are also involved in direct S1 interactions. Key residue Lys31 that is observed in both CA-INT and CA/SC-INT is the residue in common with the findings from the residue-level RMSD comparison between ACE2 crystal structures.

## Application of LDA classification vectors to ligand-bound ACE2 systems

Several recent computational studies proposed that some potent molecules either directly bind to the interface,^41–53^ to a distant site from the interface,^54^ or to the active site of ACE2^55,56^ with the goal of allosterically affecting the interactions between ACE2 and S1 and help prevent the viral entry of SARS-CoV-2. As mentioned earlier, DESRES also shared several MD simulations that were performed for ACE2 in complex with 78 compounds predicted to bind tightly to ACE2.^31^ The binding sites for these compounds are shown in Fig. 4A and grouped as follows: near S1-interacting site^6,9^ (green), inside the deep channel of ACE2 (purple), on the backside of the top part of ACE2 (orange), near the ACE2-homodimerization site^6^ (blue), and near loop that connects the beta sheets in subdomain I (cyan). The identities of these compounds are also given in Table S1.

**Figure 4:**
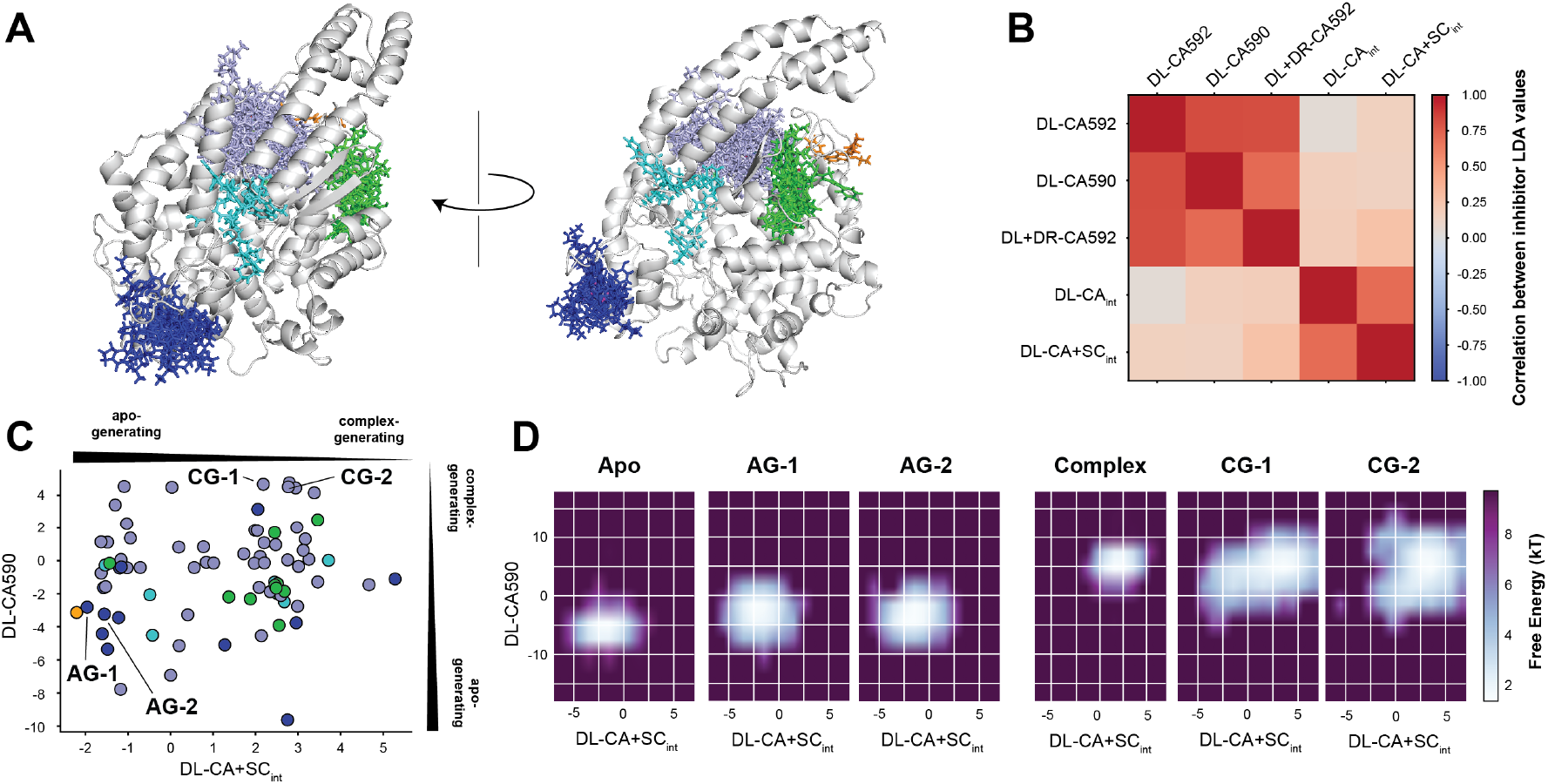
Analysis of ligand-induced conformational change in ACE2. (A) Binding sites of 78 ACE2-binding ligands simulated by D.E. Shaw Research are shown. Ligands are colored according to their binding site as described in the text. (B) The correlation matrix of different LDA predictors is shown using a colored matrix. The mean LDA projection value was determined for each of the 78 ligand-bound trajectories. Each square shows the correlation between the sets of mean values determined by a given pair of LDA predictors (red = high positive correlation, white = no correlation, blue = high negative correlation). (C) A scatter plot is shown comparing the mean LDA projection values for the CA-590 and CA/SC-INT predictors for each of the 78 ligand-bound trajectories. Each point is colored according to its binding site in panel A. (D) 2D free energy plots are projected onto CA-590 and CA/SC-INT for different trajectories. From left to right these are: i) 15 apo trajectories (100 ns each); ii) DESRES 2 *µ*s trajectory for AG-1 (CHEMBL71263), the top apo-generating ligand; iii) DESRES 2 *µ*s trajectory for AG-2 (CHEMBL2218894), the second-most apo-generating ligand; iv) 15 complex trajectories (100 ns each); v) DESRES 2 *µ*s trajectory for CG-1 (CHEMBL2105737), the top complex-generating ligand; vi) DESRES 2 *µ*s trajectory for CG-2 (CHEMBL3218576), the second-most complex-generating ligand.

In this section, we aim to predict if any of these compounds might prevent S1 binding using only the atomic positions of ACE2 structures generated in the ligand-bound ACE2 DESRES simulations. Here, we use the LDA predictors generated in the previous section, which well separate the apo and complex ACE2 structures, to classify 78 ACE2-binding compounds^31^ and investigate the extent to which these compounds are apo-generating or complex-generating, *e*.*g*. whether they produce more apo-or complex-like ACE2 conformations, respectively. We first calculate the correlation between the LDA predictors using the mean projection score values of each of the 78 ligand molecules. Fig. 4B) shows the correlation matrix for five LDA predictors (CA-592, CA-590, CA-592-DR, CA-INT and CA/SC-INT). Here, positive correlations are denoted as red whereas blue is for negative. Interestingly we observe two sets of correlated predictors: i) CA-592, CA-590, CA-592-DR, and ii) CA-INT and CA/SC-INT. Little correlation is observed between the predictors in each set. It is particularly significant that predictions from pairs such as CA-592:CA-590 and CA-INT:CA/SC-INT, show a strong correlation, as these vectors are significantly different when viewed on the ACE2 structure.

We now focus on both the CA-590 and CA/SC-INT predictors, as these yield orthogonal information about the ligand-induced ACE2 conformations, neither of which is dominated by the Pro146 conformation state. Fig. 4C compares the mean projection values obtained from CA-590 and CA/SC-INT predictors for 78 compounds. Fig. 4C shows whether the inhibitors are more apo-generating or complex-generating along the CA-590 and CA/SC-INT axes, where each ligand point is colored based on its binding site. Dual apo-generating compounds that lie in the bottom left corner of this plot are of great interest and should be further investigated to examine their ability to prevent S1-binding. In general, this scatter plot shows that there is no distinct clustering of compounds that are found in the same binding site. However, notably, the compounds near the S1-binding interface (green, except one) are clustered in the complex-complex quadrant (top right) whereas 8 of the homodimerization-site compounds (blue) that bind near dimerization loop are in the apo-apo quadrant. We ranked the compounds according to the combined number of frames predicted as “apo” and examine the top two apo-generating compounds (Table 3. To identify the most complex-generating compounds, we also take into account the proximity of the mean LDA scores to that of the complex ACE2 simulations. Thus, we picked compounds with a high combined number of frames predicted as “complex” that are near right upper edge of the complex-complex quadrant in Fig. 4C. The top two apo-generating compounds are from the homodimerization site (blue) set whereas the top two complex-generating compounds are from the deep channel (purple) set as marked on Fig. 4C. In Fig. 4D, CA-590 projection values are plotted against CA/SC-INT values as free energy maps for the apo- and complex-generating compounds. Free energy maps of apo and complex ACE2 are also provided for comparison. The apo-generating compounds have relatively similar free energy distributions compared to those of the apo whereas the complex-generating compounds have larger distributions even than that of the complex, which is particularly significant as the S1 protein was not present during any of the inhibitor-bound simulations.

**Table 3:**
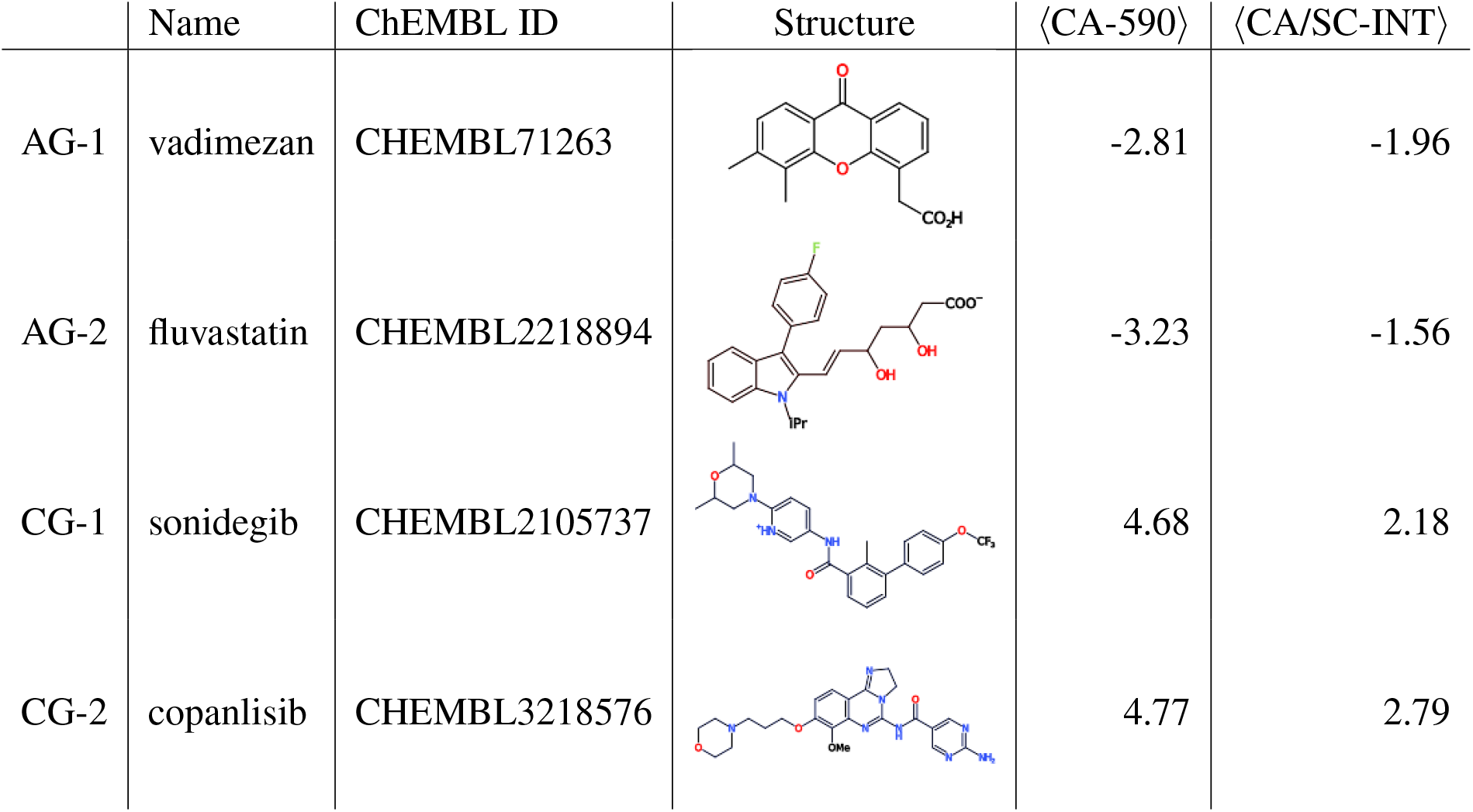
The most apo-generating and the most complex-generating compounds. Compounds were ranked according to the number of apo predicted frames using the CA-590 and CA/SC-INT LDA predictors. ⟨CA-590⟩ and ⟨CA/SC-INT⟩ show the average value of the LDA predictor across all frames in the trajectory, which are also plotted in Fig. 4C. 2-D chemical structures are generated using CDK Depict.^57^

| | Name | ChEMBL ID | Structure | $\langle \text{CA-590} \rangle$ | $\langle \text{CA/SC-INT} \rangle$ |
| --- | --- | --- | --- | --- | --- |
| AG-1 | vadimezan | CHEMBL71263 |  | -2.81 | -1.96 |
| AG-2 | fluvastatin | CHEMBL2218894 |  | -3.23 | -1.56 |
| CG-1 | sonidegib | CHEMBL2105737 |  | 4.68 | 2.18 |
| CG-2 | copanlisib | CHEMBL3218576 |  | 4.77 | 2.79 |

**Table 4:** Expectation values of contact formation between residue pairs.

| Interacting residues |  | Apo-gen. | Complex-gen. |
| --- | --- | --- | --- |
| Asp38 | Lys353 | $0.749 \pm 0.0087$ | $0.082 \pm 0.0049$ |
| Tyr41 | Asp355 | $0.710 \pm 0.0094$ | $0.111 \pm 0.0032$ |
| Tyr41 | Arg357 | $0.602 \pm 0.0091$ | $0.019 \pm 0.0021$ |
| Gln175 | Ile126 | $0.141 \pm 0.0052$ | $0.861 \pm 0.0019$ |

### Comparison of contact networks for apo- and complex-generating ligands

Contact networks are one of the ways to explore the formation and/or loss of amino acid contacts within different conformational ensembles.^58–60^ Here we calculate the distance between the closest heavy atoms of each pair of residues for all frames of a given trajectory using the MDTraj^61^ package. Note that for a residue *i*, the contact pairs (*i, i* + 1) and (*i, i* + 2) are excluded in the calculation. For all other pairs of residues we calculate *c*_*ij*_ = 1*/*1 + exp(*a*(*d*_*ij*_ −*d*_0_)), where *d*_*ij*_ is the closest distance between residues *i* and *j, d*_0_ is a cutoff equal to 5.0 Å, and *a* is a parameter controlling the steepness of the cutoff, set here to 17 Å^−1^. To estimate uncertainties in *c*_*ij*_, we divide the trajectory into four subsets and determine the average of *c*_*ij*_ for all residue pairs in each subset. We then identify significant differences between the apo- and complex-generating ligands by comparing differences between the averages with the standard deviations across the subsets. Fig. 5 shows the most significant differences in the contact networks of the trajectories for the most apo- and complex-generating ligands. These are visualized on the ACE2 structure, where blue indicates a contact that is uniquely observed in the apo-generating ligand trajectory and red in the complex-generating ligand trajectory.

We observe that many contacts are significantly different between the apo- and complex-generating ligand trajectories, which are distributed throughout the ACE2 structure. In both cases, these contact changes exist far from binding sites of each compound. The most significant contact differences are: the interactions in the S1 binding interface (black box in Fig. 5), near the dimerization loop (yellow box), and between the central helices of ACE2 (blue box). At the S1 binding interface, the salt bridge between Asp38 and Lys353 is present in the apo-generating lig- and trajectory (*c*_38−353_ = 172 *±* 16.15), but not when the complex-generating compound is bound (*c*_38−353_ = 20.75 *±* 8.69), where the uncertainties are calculated using bootstrapping as discussed above (see Table 4). Tyr41, another S1-interacting residue, loses its contacts with Asp355 and Arg357 in the complex-generating ligand trajectory, again in agreement with apo and complex trajectories. Note that all five of these residues interact with S1 protein. Near the dimerization loop, residue Gln175, which plays a role in ACE2 dimerization,^6^ makes a contact with Ile126 when the complex-generating compound is bound. Notably, we do not see any contact change in the active site of ACE2 in both cases. Together, these results clearly suggest that ACE2 is susceptible to large-scale ligand induced conformational changes, and that compounds bound to a variety of sites far from the S1-binding interface have the potential to modulate the binding free energy of S1 protein.

**Figure 5:**
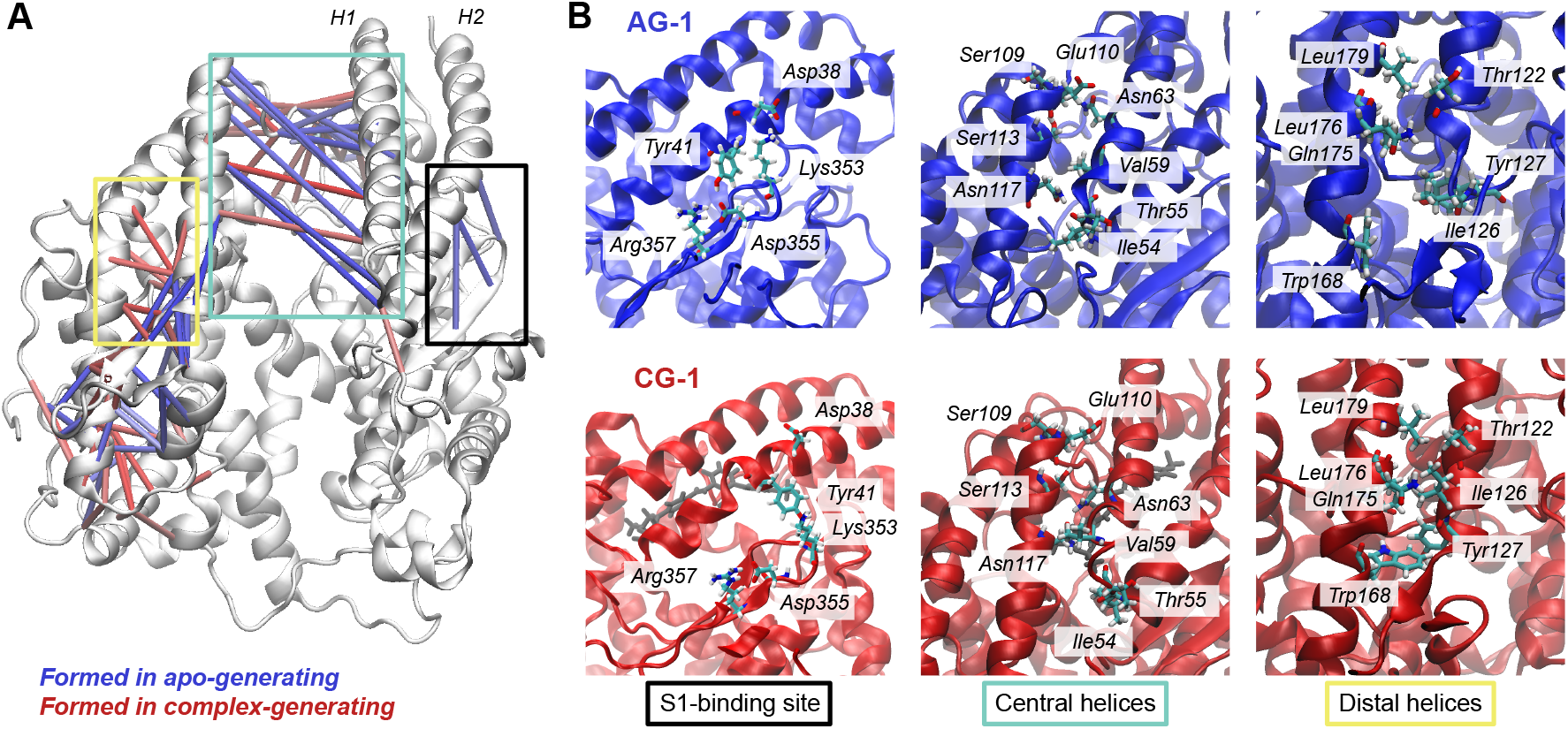
Contact networks for the most apo-(AG-1) and complex-generating (CG-1) compounds. A) Colored cylinders show contacts that are uniquely present in the simulations with
ACE2 bound to apo-generating (blue: contacts in AG-1) and complex-generating (red: contacts in CG-1) compound. B) Characteristic snapshots of the most apo- and complex-generating ligands focusing on the S1-binding site (left) the central helices (center) and distal helices (right). The corresponding regions are outlined using matching colored boxes in A.

## Discussion and Conclusions

Here we used a large set of molecular dynamics simulations of both apo and S1-bound ACE2 to identify significant structural differences that are caused by the binding of the S1 subunit of the SARS-CoV-2 tail spike. We then trained a classifier using LDA to quantitatively assess whether a given ACE2 conformation is “apo-like” or “complex-like” and applied this to a large dataset of 78 ligand-bound ACE2 trajectories made publicly available by D.E. Shaw Research.^31^ This study was enabled by the unprecedented outpouring of scientific collaboration brought about by the COVID-19 pandemic, in both the structures used for initializing the simulations,^6,36^ insight into glycosylation of both ACE2 and the S1 subunit^9–12^ and long trajectories of ACE2 used here. However, we expect that the general approach of identifying structural differences and using classifiers to prioritize compounds can be useful beyond the ACE2-S1 protein interaction. A benefit of this approach is that it explicitly prioritizes compounds that stabilize a given structural ensemble at the expense of another, which can potentially offer a means of disrupting ACE2-S1 complex formation without perturbing ACE2 function.

The simulations here were comprised only of the peptidase domain, which as shown in Fig. 1 is part of a larger protein with intracellular and transmembrane components. ACE2 can also form dimers as well as complexes with the supporting proteins TMPRSS2. Our focus on a single peptidase domain, either alone or in complex with the viral tailspike S1 domain, was motivated by the minimal contacts made across the dimer interface, and it enabled us to efficiently run longer timescale trajectories. However it is possible that ACE2 peptidase conformations could differ in the context of the full system. Interestingly, both of the most apo-generating ligands examined here (vadimezan and fluvastatin) bind near the peptidase domain dimerization loop. The potential for these compounds to alter ACE2 function by disrupting dimerization should be investigated in future studies.

One of the clearest differences between the apo and complex simulations here is the isomerization state of the Pro146 residue. This carried over from isomerization differences in the crystal structures used for the apo (1R42, cis) and complex (6M17, trans) simulations. To investigate this further we examine the Pro146 conformation in all ACE2 structures that have been published to date (Table S2). Ten of the 37 structures showed Pro146 in the trans conformation, 25 were cis, and the remaining two were intermediate between cis and trans conformations. Interestingly, both cis and trans Pro146 conformations have been shown to be compatible with S1-protein binding. The trans conformation was particularly observed for full-length ACE2 structures that are in the dimer state,^6^ as well as for an engineered trimeric form of ACE2.^62^ As Pro146 is adjacent to the dimerization loop, it is possible that isomerization is important role in function. We also note that this residue is conserved with high probability (94%) among ACE2 sequences across organisms (see ACE2 sequence alignment in Supplemental Information).

Here we examined both data generated on a GPU cluster and data generated on the Anton 2 supercomputer by D. E. Shaw Research.^30^ Anton is unique in that it can efficiently generate run very long trajectories, and those here were either 10.0 *µ*s for the apo and complex trajectories, or 2.0 *µ*s for the ligand-bound simulations. In contrast, our simulations for apo and complex ACE2 systems were performed on parallel GPUs: 15 simulations, each 100 ns in length. Longer simulations allow for new information to be revealed, as they are able to surmount large energetic barriers and explore regions of space that are farther from the initial structure. In contrast, larger ensembles of trajectories can more conclusively show whether observed differences are statistically significant; although longer trajectories can be divided into segments (or blocks) for boot-strap error analysis, this can be complicated by long correlation times and non-ergotic behavior. For this reason we consider the LDA vectors verified by independent training and testing sets to be the most reliable predictor of ACE2 structural differences.

The testing accuracy of LDA classifiers can vary significantly depending on the features used (Table 2). We find, intuitively, that it is much easier to achieve high classification accuracy on a training set than on independent testing data, although some predictors could achieve near-perfect accuracy in testing as well. The prediction accuracy goes down when training data and testing data were from different environments. Interestingly, we find there are different ways to make successful predictions that reveal complementary information (e.g. CA-590 and CA/SC-INT). While the all C-alpha approach is general and can pick up on long-range allosteric effects, the interface obviously focuses on structural information that is likely to be important for S1 binding. Here we use both and were able to find ACE2-binding ligands that were apo-generating with respect to both LDA predictors simultaneously.

Finally, note that here the LDA projections assess only the similarity of a given ligand-bound ACE2 conformational ensembles with that of the apo- and complex-ACE2 ensembles. An implicit assumption in this analysis is that higher dissimilarities between the ligand-bound and complex ensembles would result in a less-favorable binding free energy of the S1 protein. This could be directly examined in future work by calculating the free energies of binding between S1 and ACE2 restricted to different conformations. Along the same lines, to have a substantial effect, an allosteric inhibitor should have a strong binding affinity to the target. While these compounds have been shown to remain stable over long molecular dynamics simulations, binding free energies are still needed to predict whether these compounds will have allosteric effects *in vivo*.

## Methods

### Molecular Dynamics Simulations

We performed 30 independent MD simulations for apo and S1-protein bound ACE2 systems, constructed using the crystal structures 1R42.pdb^36^ and 6M17.pdb,^6^ respectively. Both structures have missing residues, therefore overlapping regions are identified and residues 21-612 are simulated in a cubic box with 10 Å of distance between the structure and the box boundaries. For ACE2, GlcNAc glycans (2-acetyl-2-deoxy-beta-d-glucosamine) are attached to six Asn residues (residue numbers are as follows: 53, 90, 103, 322, 432, 546). A single GlcNAc is attached to Asn 343 of the S1 protein. Both systems were solvated using NaCl ions at a concentration of 0.15 M. The Zn^2+^ ion at the active site is also included in the simulations. Non-bonded interactions were treated with PME and a van der Waals switching function with a cutoff of 10 Å. Simulations were run using the CHARMM36 forcefield^33–35,63^ and OpenMM version 7.5.0.^64^ We use the NPT ensemble, achieved with a Monte Carlo barostat set at 1 atm with volume moves attempted every 100 steps. The systems were equilibrated for 125 ps at 310 K using a 1.0 fs timestep. Initial velocities were given to each of the 30 replicates and ran for 100 ns at 310 K using a 2.0 fs timestep. In total, *µ*s of simulation data were collected and analyzed using MDTraj version 1.9.1.^61^.

In addition to our CHARMM MD simulations, we used simulations for the apo and S1-bound systems that were provided by D. E. Shaw Research (DESRES).^31^ All DESRES simulations were run on the Anton 2^30^ supercomputer. 10 *µ*s simulations for the apo and ACE2 were initialized from the same crystal structures used in our simulations. The Amber ff99SB-ILDN force field^65^ for proteins and the generalized Amber force field^66^ for glycosylated Asn residues were used in DESRES simulations. NAG groups (2-acetamido-2-deoxy-beta-D-glucopyranose) were attached to the same Asn residues of ACE2 and S1 as above. NaCl ions were added to neutralize the systems with a final concentration of 0.15 M. Data collection interval was set to 1.2 ns. The simulations were performed at 310 K in the NPT ensemble.

DESRES examined 5152 molecules in an FDA-investigational drug library^32^ and selected 78 out of these molecules that remained bound to ACE2 in 2 *µ*s long MD simulations. These ACE2-binding compounds are located at distinct regions on ACE2: a pocket underneath a helical bundle (residue 20-100) near top of ACE2 deep channel, a pocket involving a beta-hairpin structure (residue 346 to 360) near the S1-interacting site^6,9^ and a pocket behind a loop near the ACE2-homodimerization site^6^ (residue 131-142). Ligand-bound simulations were initiated from a different crystal structure with PDB ID: 6VW1.^9^ The same simulation parameters were used as in the previous apo and complex DESRES simulations.

### Linear Discriminant Analysis (LDA)

LDA is a machine learning method that is used for dimensionality reduction in multiclass systems. In this method, LDA builds a set of projector variables that minimize the intra-class information while maximizing the separation between classes. After LDA training, one can use these projector variables to predict classes for new, unlabeled data. In MD simulations, LDA can be used to detect differences between datasets and predict the class index of a new conformation. Classes can correspond to any labels attached to the inputs, here we use two classes for ACE2 conformations: “apo” and “complex”.

To apply LDA to an MD simulation dataset we first need to project each conformation onto a set of features (*q*_*m*_) that minimally represent all of the pertinent dynamics of the system. Here we use *x, y, z* coordinates of different groups such as C-alpha atoms and residue sidechains. The feature set *q*_*m*_ is projected onto a low-dimensional set of projector variables *y*_*m*_ using a projection matrix *W*. For the *m*-th simulation and *t*-th sample *y*_*m*_ can be written as:

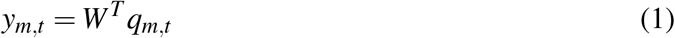

The mean of the entire dataset 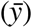 and the mean for each class 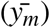 are calculated. Then, the average variance values within the classes and between the classes are computed using Eqs. 2 and 3, respectively:

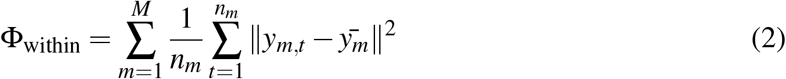

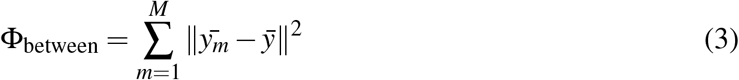

where ‖ · ‖^2^ is the Euclidean distance.

The generalized eigenvalue problem for the matrix Φ_between_/Φ_within_ was solved using singular value decomposition, as implemented in the LinearDiscriminantAnalysis class in the sci-kit learn package.^67^ After this, the first LDA mode gives the best separation between the apo and complex classes in this project. Translational and rotational degrees of freedom were removed from the LDA mode vector. The elements of this vector yields the specific residues that describe the differences between the classes and are used to draw arrows for vector visualization.

## Supporting information

Supplemental Aln File

## Acknowledgements

Authors acknowledge HPCC resources provided by the iCER at Michigan State University, and the data provided by the D. E. Shaw Research Company.

## Supporting Information Available

The following files are available free of charge.

- Filename: brief description
- Filename: brief description

This material is available free of charge via the Internet at http://pubs.acs.org/.

## Graphical TOC Entry

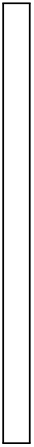

## References

(1) Samavati, L., Uhal, B. D. ACE2, Much More Than Just a Receptor for SARS-COV-2. Frontiers in Cellular and Infection Microbiology 2020, 10, 317.

(2) Zhou, P. et al. A pneumonia outbreak associated with a new coronavirus of probable bat origin. Nature 2020, 579, 270–273.

(3) Hoffmann, M., Kleine-weber, H., Schroeder, S., Mu, M. A., Drosten, C., Po, S., Hoff-mann, M., Kleine-weber, H., Schroeder, S., Kru, N. SARS-CoV-2 Cell Entry Depends on ACE2 and TMPRSS2 and Is Blocked by a Clinically Proven Article SARS-CoV-2 Cell Entry Depends on ACE2 and TMPRSS2 and Is Blocked by a Clinically Proven Protease Inhibitor. Cell 2020, 181, 271–280.

(4) Li, W., Zhang, C., Kuhn, J. H., Moore, M. J., Luo, S., Wong, S.-k., Xu, K., Vasilieva, N., Murakami, A., He, Y., Wayne, A., Guan, Y., Choe, H., Farzan, M. Receptor and viral determinants of SARS-coronavirus adaptation to human ACE2. EMBO J. 2005, 24, 1634–1643.

(5) Wu, K., Li, W., Peng, G., Li, F. Crystal structure of NL63 respiratory coronavirus receptor-binding domain complexed with its human receptor. PNAS 2009, 106, 19970–19974.

(6) Yan, R., Zhang, Y., Li, Y., Xia, L., Guo, Y., Zhou, Q. Structural basis for the recognition of SARS-CoV-2 by full-length human ACE2. Science 2020, 367, 1444–1448.

(7) Laskowski, R. A., Hutchinson, E. G., Michie, A. D., Wallace, A. C., Jones, M. L., Thorn-ton, J. M. PDBsum: A Web-based database of summaries and analyses of all PDB structures. Trends in Biochemical Sciences 1997, 22, 488–490.

(8) Laskowski, R. A., Chistyakov, V. V., Thornton, J. M. PDBsum more: New summaries and analyses of the known 3D structures of proteins and nucleic acids. Nucleic Acids Research 2005, D266–D268.

(9) Shang, J., Ye, G., Shi, K., Wan, Y., Luo, C., Aihara, H., Geng, Q., Auerbach, A., Li, F. Structural basis of receptor recognition by SARS-CoV-2. Nature 2020, 581, 221–224.

(10) Bernardi, A., Huang, Y., Harris, B., Xiong, Y., Nandi, S., Mcdonald, A., Id, R. F. Development and simulation of fully glycosylated molecular models of ACE2-Fc fusion proteins and their interaction with the SARS-CoV-2 spike protein binding domain. PLOS ONE 2020, 2, 1–12.

(11) Barros, E. P., Casalino, L., Gaieb, Z., Dommer, A. C., Wang, Y., Fallon, L., Raguette, L., Belfon, K., Simmerling, C., Amaro, R. E. The Flexibility of ACE2 in the Context of SARS-CoV-2 Infection. Biophysical Journal 2020,

(12) Chan, K. K., Dorosky, D., Sharma, P., Abbasi, S. A., Dye, J. M., Kranz, D. M., Herbert, A. S., Procko, E. Engineering human ACE2 to optimize binding to the spike protein of SARS coronavirus 2. Science 2020, 369, 1261–1265.

(13) Mehdipour, A. R., Hummer, G. Dual nature of human ACE2 glycosylation in binding to SARS-CoV-2 spike. bioRxiv 2020,

(14) Brielle, A. E. S., Schneidman-duhovny, D., Linial, M. The SARS-CoV-2 exerts a distinctive strategy for interacting with the ACE2 human receptor. Viruses 2020, 12, 497.

(15) Lupala, C. S., Li, X., Lei, J., Chen, H., Qi, J., Liu, H., Su, X.-d. Computational simulations reveal the binding dynamics between human ACE2 and the receptor binding domain of SARS-CoV-2 spike protein. bioRxiv 2020,

(16) Ali, A., Vijayan, R. Dynamics of the ACE2–SARS-CoV-2/SARS-CoV spike protein interface reveal unique mechanisms. Scientific Reports 2020, 10, 14214.

(17) Peng, C., Zhu, Z., Shi, Y., Wang, X., Mu, K., Yang, Y., Zhang, X., Xu, Z., zhu, w. Computational study of the strong binding mechanism of SARS-CoV-2 spike and ACE2. 2020.

(18) Chen, Y., Guo, Y., Pan, Y., Joe, Z. Biochemical and Biophysical Research Communications Structure analysis of the receptor binding of 2019-nCoV. Biochemical and Biophysical Research Communications 2020, 525, 135–140.

(19) Wang, Y., Liu, M., Gao, J. Enhanced receptor binding of SARS-CoV-2 through networks of hydrogen-bonding and hydrophobic interactions. Proceedings of the National Academy of Sciences 2020, 117, 13967–13974.

(20) Veeramachaneni, G. K., Thunuguntla, V. B. S. C., Bobbillapati, J., Bondili, J. S. Structural and simulation analysis of hotspot residues interactions of SARS-CoV 2 with human ACE2 receptor. Journal of Biomolecular Structure and Dynamics 2020, 1–11, PMID: 32448098.

(21) Dehury, B., Raina, V., Misra, N., Suar, M. Effect of mutation on structure, function and dynamics of receptor binding domain of human SARS-CoV-2 with host cell receptor ACE2: a molecular dynamics simulations study. Journal of Biomolecular Structure and Dynamics 2020, 1–15, PMID: 32762417.

(22) Hadi-Alijanvand, H., Rouhani, M. Studying the Effects of ACE2 Mutations on the Stability, Dynamics, and Dissociation Process of SARS-CoV-2 S1/hACE2 Complexes. Journal of Proteome Research 2020, 19, 4609–4623, PMID: 32786692.

(23) Krissinel, E., Henrick, K. Inference of Macromolecular Assemblies from Crystalline State. Journal of Molecular Biology 2007, 372, 774–797.

(24) Schymkowitz, J., Borg, J., Stricher, F., Nys, R., Rousseau, F., Serrano, L. The FoldX web server: an online force field. Nucleic acids research 2005, 33, W382–W388.

(25) Natesh, R., Schwager, S. L. U., Sturrock, E. D., Acharya, K. R. Crystal structure of the human angiotensin-converting enzyme-lisinopril complex. Nature 2003, 421, 551–554.

(26) Arndt, J. W., Hao, B., Ramakrishnan, V., Cheng, T., Chan, S. I., Chan, M. K. Crystal Structure of a Novel Carboxypeptidase from the Hyperthermophilic Archaeon Pyrococcus furiosus. Structure 2002, 10, 215–224.

(27) Brown, C. K., Madauss, K., Lian, W., Beck, M. R., Tolbert, W. D., Rodgers, D. W. Structure of neurolysin reveals a deep channel that limits substrate access. Proceedings of the National Academy of Sciences 2001, 98, 3127–3132.

(28) Uyar, A., Karamyan, V., Dickson, A. Long-Range Changes in Neurolysin Dynamics Upon Inhibitor Binding. Journal of Chemical Theory and Computation 2018, 14, 444–452.

(29) Mohammad, A., Marafie, S. K., Alshawaf, E., Abu-Farha, M., Abubaker, J., Al-Mulla, F. Structural analysis of ACE2 variant N720D demonstrates a higher binding affinity to TM-PRSS2. Life Sciences 2020, 259, 118219.

(30) Shaw, D. E. et al. Anton 2: Raising the Bar for Performance and Programmability in a Special-Purpose Molecular Dynamics Supercomputer. SC ‘14: Proceedings of the International Conference for High Performance Computing, Networking, Storage and Analysis. 2014; pp 41–53.

(31) D. E. Shaw Research, Molecular Dynamics Simulations Related to SARS-CoV-2. 2020.

(32) Mendez, D. et al. ChEMBL : towards direct deposition of bioassay data. Nucleic Acids Research 2019, 47, 930–940.

(33) Brooks, B. R., Bruccoleri, R. E., Olafson, B. D., States, D. J., Swaminathan, S., Karplus, M. CHARMM: A program for macromolecular energy, minimization, and dynamics calculations. Journal of Computational Chemistry 1983, 4, 187–217.

(34) MacKerell Jr., A. D., Brooks III, C. L., Nilsson, L., Roux, B., Won, Y., Karplus, M. CHARMM: The Energy Function and Its Parameterization with an Overview of the Pro-gram. In The Encyclopedia of Computational Chemistry; et al. Schleyer, P., Ed., John Wiley & Sons: Chichester, 1998; Vol. 1; pp 271–277.

(35) Brooks, B. R. et al. CHARMM: The biomolecular simulation program. Journal of Computational Chemistry 2009, 30, 1545–1614.

(36) Towler, P., Staker, B., Prasad, S. G., Menon, S., Tang, J., Parsons, T., Ryan, D., Fisher, M., Williams, D., Dales, N. A., Patane, M. A., Pantoliano, M. W. ACE2 X-Ray Structures Reveal a Large Hinge-bending Motion Important for Inhibitor Binding and Catalysis. J. Biol. Chem. 2004, 279, 17996–18007.

(37) Pele, O., Werman, M. A linear time histogram metric for improved sift matching. Computer Vision–ECCV 2008. 2008; pp 495–508.

(38) Pele, O., Werman, M. Fast and robust earth mover’s distances. 2009 IEEE 12th International Conference on Computer Vision. 2009; pp 460–467.

(39) Fisher, R. A. The Use of Multiple Measurements in Taxonomic Problems. Annals of Eugenics 1936, 7, 179–188.

(40) Rao, C. R. The Utilization of Multiple Measurements in Problems of Biological Classification. Journal of the Royal Statistical Society. Series B (Methodological) 1948, 10, 159–203.

(41) Smith, M., Smith, J. C. Repurposing Therapeutics for COVID-19: Supercomputer-Based Docking to the SARS-CoV-2 Viral Spike Protein and Viral Spike Protein-Human ACE2 Interface. 2020; DOI:10.26434/chemrxiv.11871402.v4.

(42) Trezza, A., Iovinelli, D., Santucci, A., Prischi, F., Spiga, O. An integrated drug repurposing strategy for the rapid identification of potential SARS - CoV - 2 viral inhibitors. Scientific Reports 2020, 1–8.

(43) Jani, V., Koulgi, S., Uppuladinne V N, M., Sonavane, U., Joshi, R. Computational Drug Repurposing Studies on the ACE2-Spike (RBD) Interface of SARS-CoV-2. 2020.

(44) Sandeep, S., McGregor, K. Energetics Based Modeling of Hydroxychloroquine and Azithromycin Binding to the SARS-CoV-2 Spike (S)Protein - ACE2 Complex. 2020.

(45) Khelfaoui, H., Harkati, D., Saleh, B. A. Molecular docking, molecular dynamics simulations and reactivity, studies on approved drugs library targeting ACE2 and SARS-CoV-2 binding with ACE2. Journal of Biomolecular Structure and Dynamics 2020, 1–17.

(46) Yang, J., Petitjean, S. J. L., Koehler, M., Zhang, Q., Dumitru, A. C., Chen, W., Derclaye, S., Vincent, S. P., Soumillion, P., Alsteens, D. Molecular interaction and inhibition of SARS-CoV-2 binding to the ACE2 receptor. Nature Communications 2020, 11, 4541.

(47) Han, Y., Kral, P. Computational Design of ACE2-Based Peptide Inhibitors of SARS-CoV-2. ACS Nano 2020, 14, 5143–5147, PMID: 32286790.

(48) Yun, Y., Song, H., Ji, Y., Huo, D., Han, F., Li, F., Jiang, N. Identification of therapeutic drugs against COVID-19 through computational investigation on drug repurposing and structural modification. Journal of Biomedical Research 2020, 34, 458–469.

(49) Prajapat, M., Shekhar, N., Sarma, P., Avti, P., Singh, S., Kaur, H., Bhattacharyya, A., Kumar, S. Journal of Molecular Graphics and Modelling Virtual screening and molecular dynamics study of approved drugs as inhibitors of spike protein S1 domain and ACE2 interaction in SARS-. Journal of Molecular Graphics and Modelling 2020, 101, 107716.

(50) Deganutti, G., Prischi, F., Reynolds, C. A. Supervised molecular dynamics for exploring the druggability of the SARS-CoV-2 spike protein. Journal of Computer-Aided Molecular Design 2020,

(51) Wang, G. et al. Dalbavancin binds ACE2 to block its interaction with SARS-CoV-2 spike protein and is effective in inhibiting SARS-CoV-2 infection in animal models. Cell Research 2021, 31, 17–24.

(52) Sekar, P. C., Rajasekaran, R. Could Dermaseptin Analogue be a Competitive Inhibitor for ACE2 Towards Binding with Viral Spike Protein Causing COVID19 ?: Computational Investigation. International Journal of Peptide Research and Therapeutics 2021, 1–14.

(53) Baby, K., Maity, S., Mehta, C. H., Suresh, A., Nayak, U. Y., Nayak, Y. SARS-CoV-2 entry inhibitors by dual targeting TMPRSS2 and ACE2: An in silico drug repurposing study. European Journal of Pharmacology 2021, 896, 173922.

(54) Gross, L. Z. F., Sacerdoti, M., Piiper, A., Leroux, A. E., Biondi, R. M. ACE2, the Receptor that Enables Infection by SARS-CoV-2: Biochemistry, Structure, Allostery and Evaluation of the Potential Development of ACE2 Modulators. ChemMedChem 2020, 15, 1682–1690.

(55) Terali, K., Baddal, B., Gülcan, H. O. Prioritizing potential ACE2 inhibitors in the COVID-19 pandemic: Insights from a molecular mechanics-assisted structure-based virtual screening experiment. Journal of molecular graphics & modelling 2020, 100, 107697.

(56) Joshi, T., Joshi, T., Sharma, P., Mathpal, S., Pundir, H. In silico screening of natural compounds against COVID-19 by targeting Mpro and ACE2 using molecular docking. Eur. Rev. Med. Pharmacol. Sci. 2020, 24, 4529–4536.

(57) CDK Depict. https://www.simolecule.com/cdkdepict/depict.html.

(58) Di Paola, L., De Ruvo, M., Paci, P., Santoni, D., Giuliani, A. Protein Contact Networks: An Emerging Paradigm in Chemistry. Chemical Reviews 2013, 113, 1598–1613, PMID: 23186336.

(59) Dokholyan, N. V. Controlling Allosteric Networks in Proteins. Chemical Reviews 2016, 116, 6463–6487, PMID: 26894745.

(60) Ji, S., Oruc, T., Mead, L., Rehman, M. F., Thomas, C. M., Butterworth, S., Winn, P. J. DeepCDpred: Inter-residue distance and contact prediction for improved prediction of protein structure. PLOS ONE 2019, 14, 1–15.

(61) McGibbon, R. T., Beauchamp, K. A., Harrigan, M. P., Klein, C., Swails, J. M., Hernán-dez, C. X., Schwantes, C. R., Wang, L.-P., Lane, T. J., Pande, V. S. MDTraj: A Modern Open Library for the Analysis of Molecular Dynamics Trajectories. Biophysical Journal 2015, 109, 1528 – 1532.

(62) Xiao, T. et al. A trimeric human angiotensin-converting enzyme 2 as an anti-SARS-CoV-2 agent in vitro. bioRxiv 2020,

(63) Huang, J., MacKerell, A. CHARMM36 all atom additive protein force field: Validation based on comparison to NMR data. Journal of Computational Chemistry 2013, 34, 2135–2145.

(64) Eastman, P., Swails, J., Chodera, J. D., McGibbon, R. T., Zhao, Y., Beauchamp, K. A., Wang, L.-P., Simmonett, A. C., Harrigan, M. P., Brooks, B. R., Pande, V. S. OpenMM 7: Rapid Development of High Performance Algorithms for Molecular Dynamics. PLoS Computational Biology 2017, 13, :e1005659.

(65) Lindorff-Larsen, K., Piana, S., Palmo, K., Maragakis, P., Klepeis, J. L., Dror, R. O., Shaw, D. E. Improved side-chain torsion potentials for the Amber ff99SB protein force field. Proteins: Structure, Function and Bioinformatics 2010, 78, 1950–1958.

(66) Wang, J., Wolf, R. M., Caldwell, J. W., Kollman, P. A., Case, D. A. Development and testing of a general amber force field. Journal of Computational Chemistry 2004, 25, 1157–1174.

(67) Pedregosa, F. et al. Scikit-learn: Machine Learning in Python. Journal of Machine Learning Research 2011, 12, 2825–2830.

