## Supplemental Information for "Perturbation of ACE2 structural ensembles by SARS-CoV-2 spike protein binding"

March 2, 2021

Table S1: Chemical structure of ACE2 ligands simulated in the D. E. Shaw Research dataset [1]. Note that ligands doxazosin and pimecrolimus each had two trajectories, starting from different binding sites.

| Name | ChEMBL ID | Structure | Av. LDA (CA-590) | Av. LDA (CA/SC-INT) | Binding site |
|---|---|---|---|---|---|
| acalabrutinib | CHEMBL3707348 | 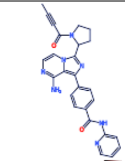 | -0.14 | 1.01 | 3 |
| aclidinium | CHEMBL1194325 | 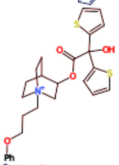 | 2.04 | 2.97 | 3 |
| afeletecan | CHEMBL134380 | 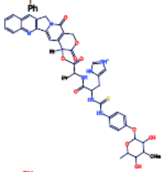 | 0.04 | 2.75 | 3 |
| amentoflavone | CHEMBL63354 | 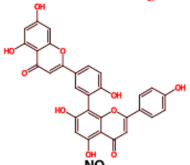 | -4.42 | -1.60 | 5 |
| azathioprine | CHEMBL1200400 | 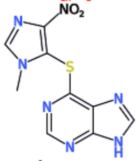 | 2.26 | -1.03 | 3 |
| barusiban | CHEMBL2218898 | 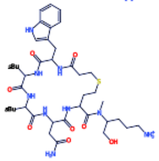 | -0.34 | 3.01 | 3 |

| | | | | | |
|---|---|---|---|---|---|
| bekanamycin | CHEMBL1237048 | 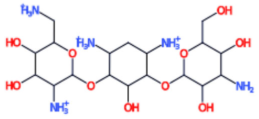 | 4.52 | 2.77 | 3 |
| bendamustine | CHEMBL1201734 | 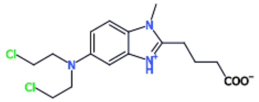 | -2.06 | -0.49 | 4 |
| berzosertib | CHEMBL3989870 | 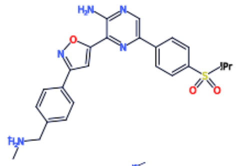 | -0.42 | 0.56 | 3 |
| birinapant | CHEMBL3039522 | 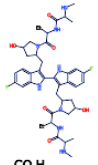 | -7.77 | -1.17 | 3 |
| brequinar | CHEMBL300058 | 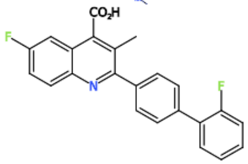 | -0.15 | -1.43 | 2 |
| brigatinib | CHEMBL3545311 | 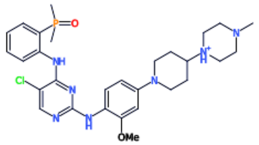 | 0.42 | 1.72 | 3 |
| bromocriptine | CHEMBL1200503 | 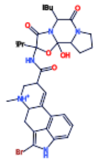 | -3.44 | -1.22 | 5 |
| brompheniramine | CHEMBL1200961 | 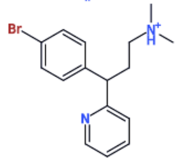 | -5.35 | -1.48 | 5 |

| | | | | | |
|---|---|---|---|---|---|
| bunazosin | CHEMBL188185 | 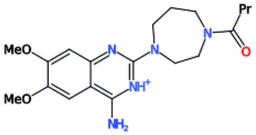 | 2.51 | 3.46 | 2 |
| capreomycin | CHEMBL1201252 | 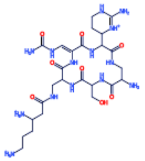 | -0.43 | -1.02 | 3 |
| ceritinib | CHEMBL2403108 | 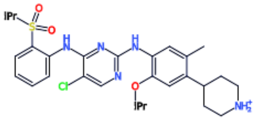 | -2.76 | 2.99 | 3 |
| chloroquine | CHEMBL1326 | 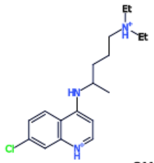 | 3.42 | -1.30 | 3 |
| cimicoxib | CHEMBL435381 | 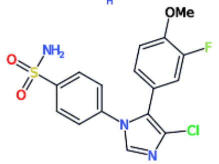 | -1.66 | 2.48 | 2 |
| conivaptan | CHEMBL1201108 | 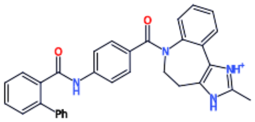 | 0.02 | 3.71 | 4 |
| copanlisib | CHEMBL3218576 | 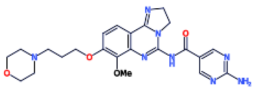 | 4.77 | 2.79 | 3 |
| dabigatran | CHEMBL48361 | 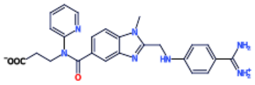 | -1.28 | 3.46 | 3 |

| | | | | | |
|---|---|---|---|---|---|
| dacomitinib | CHEMBL2105719 | 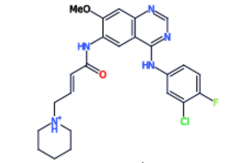 | 4.49 | 0.03 | 3 |
| dasatinib | CHEMBL1421 | 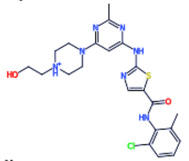 | -1.62 | 2.08 | 3 |
| delavirdine | CHEMBL593 | 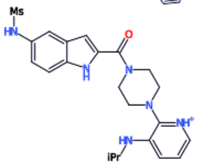 | -1.96 | 2.50 | 2 |
| desvenlafaxine | CHEMBL1118 | 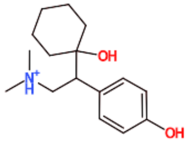 | -0.27 | -1.54 | 4 |
| deutivacaftor | CHEMBL2010601 | 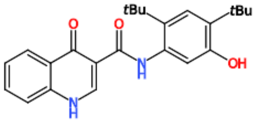 | 1.13 | -1.48 | 3 |
| dimenhydrinate | CHEMBL1200406 | 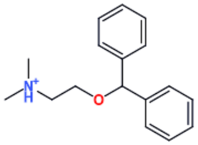 | -6.92 | 0.01 | 3 |
| disopyramide | CHEMBL1201020 | 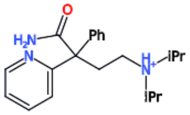 | 0.07 | -1.16 | 3 |
| doxazosin | CHEMBL1200561 | 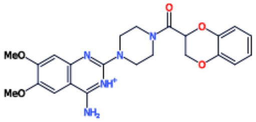 | -1.86 | 2.68 | 2 |

| | | | | | |
|---|---|---|---|---|---|
| doxazosin | CHEMBL1200561 | 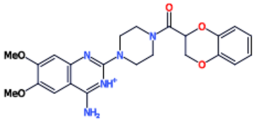 | 1.86 | 1.98 | 3 |
| e7107 | CHEMBL4297278 | 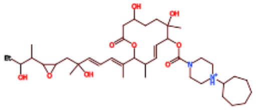 | 0.09 | 3.25 | 3 |
| elamipretide | CHEMBL3833370 | 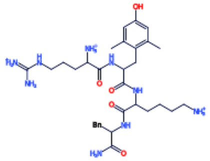 | 0.84 | 0.78 | 3 |
| elbasvir | CHEMBL3039514 | 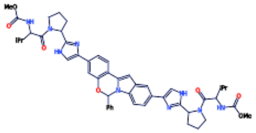 | -1.87 | 2.34 | 3 |
| eletriptan | CHEMBL1201003 | 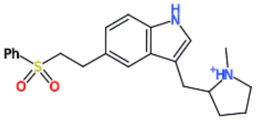 | 1.32 | 3.17 | 3 |
| elvitegravir | CHEMBL204656 | 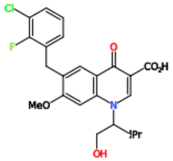 | -2.31 | 1.88 | 2 |
| fluoxetine | CHEMBL1201082 | 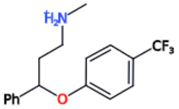 | -0.45 | -1.20 | 5 |
| fluvastatin | CHEMBL2218894 | 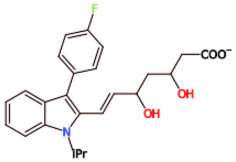 | -3.23 | -1.56 | 5 |

| | | | | | |
|---|---|---|---|---|---|
| gemifloxacin | CHEMBL1200621 | 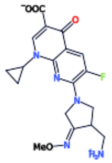 | -0.37 | -1.16 | 5 |
| gentamicin | CHEMBL329592 | 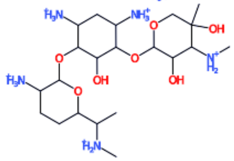 | -2.11 | 2.60 | 3 |
| imatinib | CHEMBL1642 | 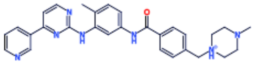 | 1.09 | 2.20 | 3 |
| ipamorelin | CHEMBL58547 | 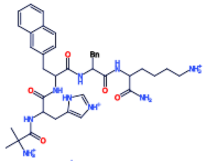 | -0.13 | 2.21 | 3 |
| kanamycin | CHEMBL1384 | 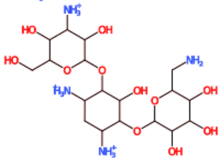 | 0.25 | 2.11 | 3 |
| KPT-9274 | CHEMBL4297467 | 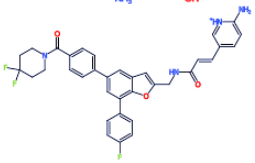 | -0.15 | 1.97 | 3 |
| lixivaptan | CHEMBL49429 | 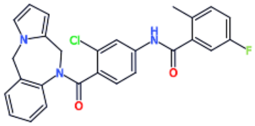 | -3.91 | 2.55 | 2 |
| mefloquine | CHEMBL416956 | 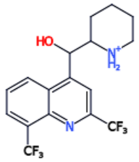 | -9.62 | 2.75 | 5 |

| | | | | | |
|---|---|---|---|---|---|
| methylnaltrexone | CHEMBL1186579 | 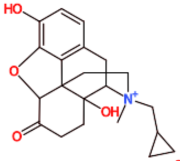 | -2.49 | 2.68 | 4 |
| methylprednisolone | CHEMBL190259 | 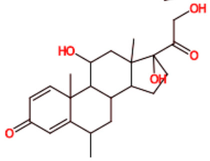 | -1.11 | 5.28 | 5 |
| metronidazole | CHEMBL1200869 | 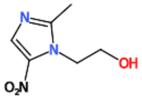 | 4.54 | -1.09 | 3 |
| midazolam | CHEMBL1200420 | 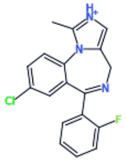 | -5.09 | 1.28 | 5 |
| modithromycin | CHEMBL1673372 | 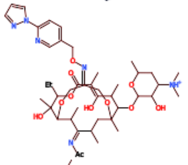 | -0.09 | 0.84 | 3 |
| nafithromycin | CHEMBL4297519 | 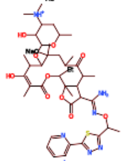 | 4.45 | 2.94 | 3 |
| netilmicin | CHEMBL1551235 | 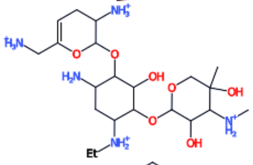 | 4.16 | 3.38 | 3 |
| netupitant | CHEMBL206253 | 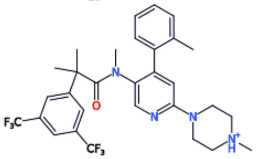 | -4.53 | 2.14 | 3 |

| | | | | | |
|---|---|---|---|---|---|
| nevirapine | CHEMBL57 | 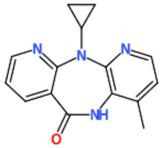 | -1.45 | 4.67 | 3 |
| orphenadrine | CHEMBL1200395 | 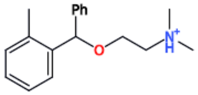 | -3.27 | 0.41 | 3 |
| panobinostat | CHEMBL3545368 | 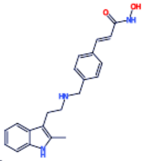 | 3.16 | 2.05 | 5 |
| pelitinib | CHEMBL607707 | 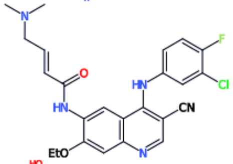 | -3.13 | -2.21 | 1 |
| perphenazine | CHEMBL567 | 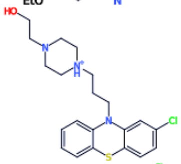 | -1.49 | 2.46 | 2 |
| pimecrolimus | CHEMBL1200686 | 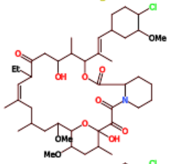 | -3.76 | 2.96 | 5 |
| pimecrolimus | CHEMBL1200686 | 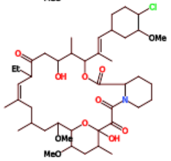 | 0.82 | 0.19 | 3 |
| pioglitazone | CHEMBL1715 | 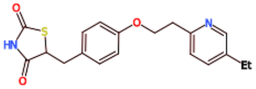 | 1.37 | -0.94 | 3 |

| | | | | | |
|---|---|---|---|---|---|
| pramoxine | CHEMBL1198 | 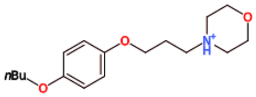 | -0.41 | -0.71 | 3 |
| prazosin | CHEMBL1347191 | 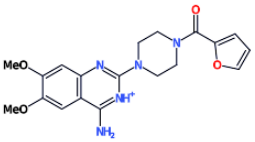 | 1.71 | 2.45 | 2 |
| primaquine | CHEMBL3217110 | 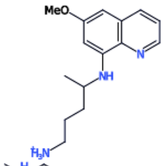 | -1.30 | 2.45 | 4 |
| prochlorperazine | CHEMBL1200587 | 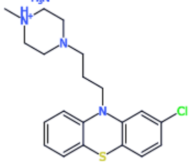 | -1.63 | -1.57 | 5 |
| propylhexedrine | CHEMBL2105275 | 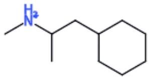 | -4.50 | -0.42 | 4 |
| rabeprazole | CHEMBL1200930 | 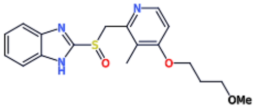 | -5.14 | 0.20 | 3 |
| ruzasvir | CHEMBL3833385 | 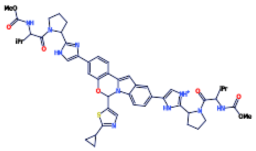 | -0.75 | -1.63 | 3 |
| sivifene | CHEMBL1628305 | 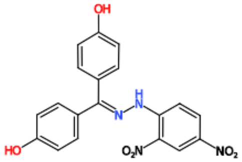 | -1.38 | 2.51 | 2 |

| | | | | | |
|---|---|---|---|---|---|
| sonidegib | CHEMBL2105737 | 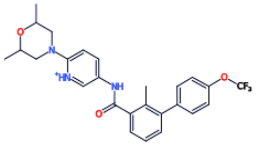 | 4.68 | 2.18 | 3 |
| ticlopidine | CHEMBL1717 | 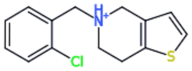 | 1.15 | -1.63 | 3 |
| tivozanib | CHEMBL1289494 | 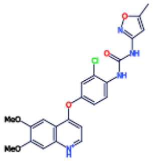 | -2.19 | 1.37 | 2 |
| tobramycin | CHEMBL1200780 | 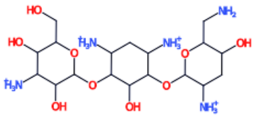 | 1.85 | 2.04 | 3 |
| topiramate | CHEMBL220492 | 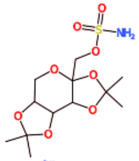 | -1.57 | -1.53 | 3 |
| TT-232 | CHEMBL539934 | 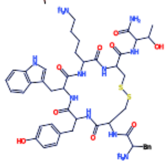 | 0.97 | 2.47 | 3 |
| vadimezan | CHEMBL71263 | 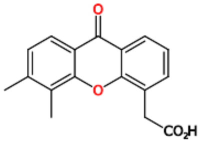 | -2.81 | -1.96 | 5 |

| | | | | | |
|---|---|---|---|---|---|
| ziritaxestat | CHEMBL3828074 | 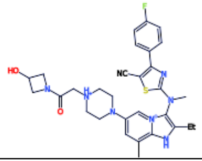 | 0.63 | 3.12 | 3 |

Table S2: Isomerization states of PRO145 of ACE2 for solved structure in the literature. Isomerization states were determined using the corresponding $\omega$ backbone angle, with a cutoff of $0.5$ radians from the basins defining cis $(0.0)$ and trans $(\pm\pi)$. Angles outside of this range are denoted “int” for intermediate. Note that trimer oligomerization states are engineered trimers of ACE2 protease domains [2].

| PDB ID | Iso. state | Olig. state | Spike bound | Method |
|---|---|---|---|---|
| 1r42 | cis | monomer | apo | X-ray |
| 1r4l | cis | monomer | apo | X-ray |
| 2ajf | cis | monomer | apo | X-ray |
| 3d0g | cis | monomer | SARS-CoV | X-ray |
| 3d0h | cis | monomer | SARS-CoV | X-ray |
| 3d0i | cis | monomer | SARS-CoV | X-ray |
| 3kbh | int | monomer | NL63 | X-ray |
| 3sci | cis | monomer | SARS-CoV | X-ray |
| 3scj | cis | monomer | SARS-CoV | X-ray |
| 3sck | cis | monomer | SARS-CoV | X-ray |
| 3scl | cis | monomer | SARS-CoV | X-ray |
| 6acg | cis | monomer | SARS-CoV | Cryo-EM |
| 6acj | cis | monomer | SARS-CoV | Cryo-EM |
| 6ack | cis | monomer | SARS-CoV | Cryo-EM |
| 6cs2 | cis | monomer | SARS-CoV | Cryo-EM |
| 6lzg | cis | monomer | SARS-CoV-2 | X-ray |
| 6m0j | cis | monomer | SARS-CoV-2 | X-ray |
| 6m17 | trans | dimer | SARS-CoV-2 | Cryo-EM |
| 6m18 | int | dimer | apo | Cryo-EM |
| 6m1d | trans | dimer | SARS-CoV-2 | Cryo-EM |
| 6vw1 | cis | monomer | SARS-CoV-2 | X-ray |
| 7a91 | cis | monomer | SARS-CoV-2 | Cryo-EM |
| 7a92 | cis | monomer | SARS-CoV-2 | Cryo-EM |
| 7a94 | cis | monomer | SARS-CoV-2 | Cryo-EM |
| 7a95 | cis | monomer | SARS-CoV-2 | Cryo-EM |
| 7a96 | cis | monomer | SARS-CoV-2 | Cryo-EM |
| 7a97 | cis | monomer | SARS-CoV-2 | Cryo-EM |
| 7a98 | cis | monomer | SARS-CoV-2 | Cryo-EM |
| 7c8d | cis | monomer | SARS-CoV-2 | Cryo-EM |
| 7ct5 | trans | trimer | SARS-CoV-2 | Cryo-EM |
| 7kj2 | trans | trimer | SARS-CoV-2 | Cryo-EM |
| 7kj3 | trans | trimer | SARS-CoV-2 | Cryo-EM |
| 7kj4 | trans | trimer | SARS-CoV-2 | Cryo-EM |
| 7kmb | trans | monomer | SARS-CoV-2 | Cryo-EM |
| 7kms | trans | monomer | SARS-CoV-2 | Cryo-EM |
| 7kmz | trans | monomer | SARS-CoV-2 | Cryo-EM |
| 7knb | trans | monomer | SARS-CoV-2 | Cryo-EM |

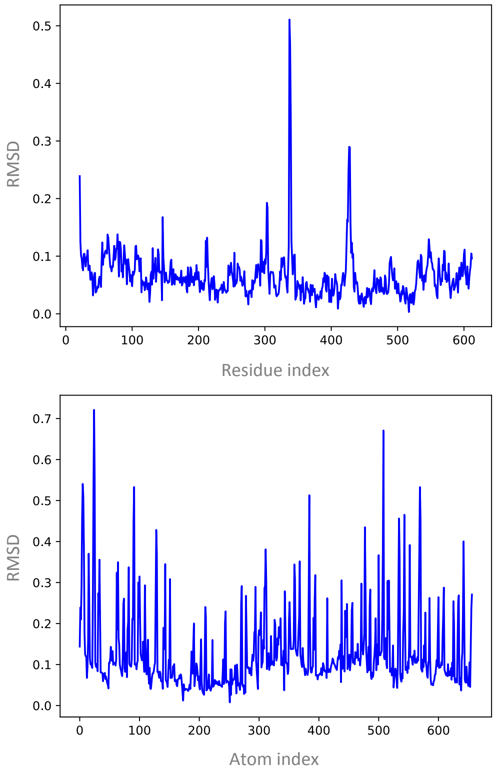

Figure S1: C$\alpha$ **Residue-level RMSD profiles between the apo and the complex ACE2 structures (top). All-atom RMSD for atoms in N-terminus helices (H1 and H2: residues 21-100) (bottom).**

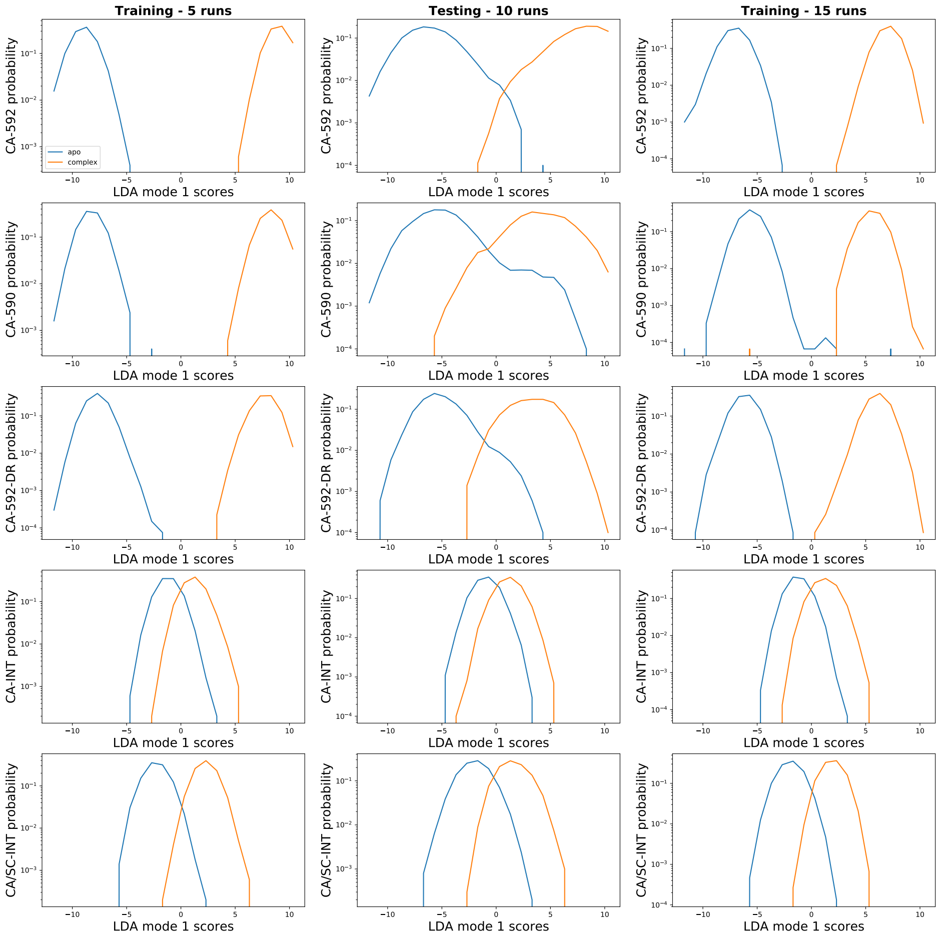

Figure S2: **Probability distributions of LDA projections. Training sets are randomly chosen from three subsets of five runs.**

### References

[1] D. E. Shaw Research. Molecular Dynamics Simulations Related to SARS-CoV-2, 2020.

[2] Tianshu Xiao, Jianming Lu, Jun Zhang, Rebecca I. Johnson, Lindsay G.A. McKay, Nadia Storm, Christy L. Lavine, Hanqin Peng, Yongfei Cai, Sophia Rits-Volloch, Shen Lu, Brian D. Quinlan, Michael Farzan, Michael S. Seaman, Anthony Griffiths, and Bing Chen. A trimeric human angiotensin-converting enzyme 2 as an anti-sars-cov-2 agent in vitro. *bioRxiv*, 2020.

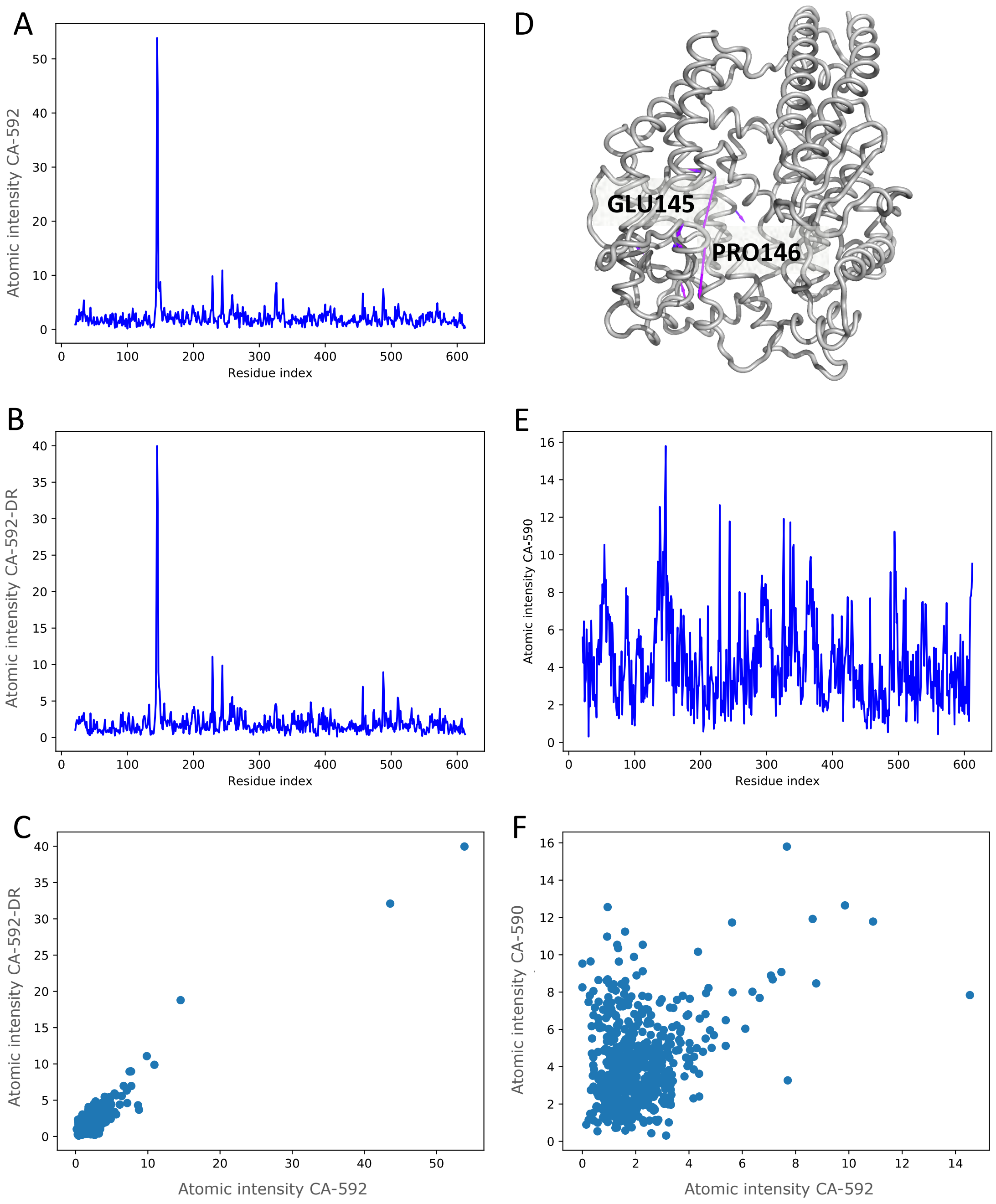

Figure S3: **(A) Atomic intensity values of LDA CA-592. (B) Atomic intensity values of LDA CA-592-DR. (C) Comparison of atomic intensity values between CA-592 and CA-592-DR. (D) Mode vector representation of LDA CA-592-DR. (E) Atomic intensity values of LDA CA-590. (F) Comparison of atomic intensity values between CA-592 and CA-590.**
